# Loss of Nuclear DNA Ligase III Reverts PARP Inhibitor Resistance in BRCA1/53BP1 Double-deficient Cells by Exposing ssDNA Gaps

**DOI:** 10.1101/2021.03.24.436323

**Authors:** Mariana Paes Dias, Vivek Tripathi, Ingrid van der Heijden, Ke Cong, Eleni-Maria Manolika, Jinhyuk Bhin, Ewa Gogola, Panagiotis Galanos, Stefano Annunziato, Cor Lieftink, Miguel Andújar-Sánchez, Sanjiban Chakrabarty, Graeme C. M. Smith, Marieke van de Ven, Roderick L. Beijersbergen, Jirina Bartkova, Sven Rottenberg, Sharon Cantor, Jiri Bartek, Arnab Ray Chaudhuri, Jos Jonkers

## Abstract

Inhibitors of poly(ADP-ribose) (PAR) polymerase (PARPi) have entered the clinic for the treatment of homologous recombination (HR)-deficient cancers. Despite the success of this approach, preclinical and clinical research with PARPi has revealed multiple resistance mechanisms, highlighting the need for identification of novel functional biomarkers and combination treatment strategies. Functional genetic screens performed in cells and organoids that acquired resistance to PARPi by loss of 53BP1 identified loss of LIG3 as an enhancer of PARPi toxicity in BRCA1-deficient cells. Enhancement of PARPi toxicity by LIG3 depletion is dependent on BRCA1 deficiency but independent of the loss of 53BP1 pathway. Mechanistically, we show that LIG3 loss promotes formation of MRE11-mediated post-replicative ssDNA gaps in BRCA1-deficient and BRCA1/53BP1 double-deficient cells exposed to PARPi, leading to an accumulation of chromosomal abnormalities. LIG3 depletion also enhances efficacy of PARPi against BRCA1-deficient mammary tumors in mice, suggesting LIG3 as a potential therapeutic target.

## INTRODUCTION

Defects in DNA repair result in genome instability and thereby contribute to the development and progression of cancer. Alterations in high-fidelity DNA repair genes lead to a greater reliance on compensatory error-prone repair pathways for cellular survival. This does not only result in the accumulation of tumor-promoting mutations, but also provides cancer-specific vulnerabilities that can be exploited for targeted cancer therapy. The first example of such targeted approach was the use of poly(ADP-ribose) polymerase (PARP) inhibitors (PARPi) in the treatment of *BRCA1* or *BRCA2* deficient tumors defective in the error-free repair of DNA double-strand breaks (DSBs) through homologous recombination (HR) (Bryant *et al*., 2005; Farmer *et al*., 2005).

PARP1, which is the main target for PARPi is involved in various cellular processes, including the sensing of DNA single-strand breaks (SSBs), repair of DNA DSBs, stabilization of replication forks (RFs), chromatin remodeling (reviewed by Ray Chaudhuri and Nussenzweig 2017) and the sensing of unligated Okazaki fragments during DNA replication (Hanzlikova *et al*., 2018). Upon DNA damage, PARP1 is rapidly recruited to sites of DNA damage where it post-translationally modifies substrate proteins by synthesizing poly(ADP-ribose) (PAR) chains in a process known as poly(ADP-ribosyl)ation (PARylation). During this process, PARP1 itself is a target of PARylation and the resulting PAR chains serve as a platform for the recruitment of downstream repair factors. AutoPARylation of PARP1 also enhances its release from DNA, which is essential for various DNA repair processes (Pascal and Ellenberger, 2015).

Initially, it was proposed that PARPi act through catalytic inhibition of PARP1, which prevents efficient repair of SSBs resulting in RF collapse and subsequent generation of DSBs during DNA replication (Lupo and Trusolino, 2014). However, later studies have demonstrated that several PARPi also trap PARP1 onto chromatin, resulting in the collapse of RFs that hit trapped PARP1 (Helleday, 2011; Murai *et al*., 2012, 2014). PARPi-treated *BRCA1/2*-defective cells can only employ error-prone repair to resolve the DSBs caused by RF collapse, resulting in accumulation of chromosomal aberrations and cell death by mitotic catastrophe (Lupo and Trusolino, 2014). Successful clinical trials have resulted in the recent approval of different PARPi for treatment of patients with *BRCA1/2*-mutant ovarian and breast cancers (Pilié *et al*., 2019). Moreover, antitumor activity of PARPi has been observed across multiple other cancer types, such as prostate and gastrointestinal cancers (Pilié *et al*., 2019).

Despite the success of this approach, multiple mechanisms of resistance to PARPi have been identified. Preclinical studies have shown that PARPi resistance can be induced by upregulation of the P-glycoprotein drug efflux transporter (Evers *et al*., 2008; Rottenberg *et al*., 2008), PARP1 downregulation/inactivation (Murai *et al*., 2012; Pettitt *et al*., 2013), mutations that abolish PARP1 trapping (Pettitt *et al*., 2018), and loss of the PAR glycohydrolase (PARG) responsible for PAR degradation (Pascal and Ellenberger, 2015; Gogola *et al*., 2018). Sensitivity to PARPi resistance may also be reduced by mechanisms that restore RF protection in the absence of BRCA1/2 (Ray Chaudhuri *et al*., 2016; Rondinelli *et al*., 2017; Lee *et al*., 2018).

The best-studied mechanisms of PARPi resistance in BRCA1/2-deficient cells involve restoration of HR activity via re-activation of BRCA1/2 function or via loss of factors that govern DSB end-protection in BRCA1-deficient cells. HR restoration due to re-established BRCA1/2 function has been observed in patients with PARPi-resistant breast cancer (Barber *et al*., 2013; Afghahi *et al*., 2017) and ovarian cancer (Edwards *et al*., 2008; Barber *et al*., 2013; Kondrashova *et al*., 2017). Restoration of HR via loss of DSB end-protection in *BRCA1*-associated tumors may be achieved by loss of 53BP1, RIF1, REV7, or components of the shieldin complex and the CST complex (Bouwman *et al*., 2010; Bunting *et al*., 2010; Zimmermann *et al*., 2013; Chapman *et al*., 2013; Escribano-Díaz *et al*., 2013; Feng *et al*., 2013; Jaspers *et al*., 2013; Boersma *et al*., 2015; Xu *et al*., 2015; Noordermeer *et al*., 2018; Dev *et al*., 2018; Ghezraoui *et al*., 2018; Gupta *et al*., 2018). Altogether, these studies underscore the high selective pressure for PARPi-treated tumors to restore HR for survival.

Drug resistance often comes at a fitness cost due to collateral vulnerabilities which can be exploited to improve therapy response. PARG inactivation causes PARPi resistance but results in increased sensitivity to ionizing radiation (IR) and temozolomide (Amé *et al*., 2009; Gogola *et al*., 2018). BRCA1-deficient tumors that acquired resistance to PARPi due to loss of the 53BP1 pathway have also been shown to become more radiosensitive (Barazas *et al*., 2019). In a similar fashion, loss of the NHEJ factors LIG4 or XRCC4 results in resistance to the DNA-damaging agent topotecan in ATM-deficient cells at the cost of increased radiosensitivity (Balmus *et al*., 2019). However, not much is known about the vulnerabilities that can be exploited to re-sensitize BRCA1- deficient PARPi resistant tumors to PARPi treatments again. In this study, we identified DNA ligase III (LIG3), a known SSB and DSB repair factor (Caldecott *et al*., 1996; Cappelli *et al*., 1997; Wang *et al*., 2005; Simsek *et al*., 2011), as a collateral vulnerability of BRCA1-deficient cells with acquired PARPi resistance due to loss of DSB end-protection. We further show that loss of LIG3 enhances the toxicity of PARPi in these cells and dissect the mechanisms that render LIG3 as a potential therapeutic target to overcome PARPi resistance.

## RESULTS

### Functional Genetic Dropout Screens Identify LIG3 as a Modulator of PARPi-resistance in BRCA1/53BP1 Double-deficient Cells

To identify acquired vulnerabilities in BRCA1-deficient cells which developed PARPi resistance via BRCA1-independent restoration of HR, we carried out functional genetic dropout screens in two types of cellular models deficient for BRCA1, p53 and 53BP1. The first screen was performed in genetically well-defined *Brca1^−/−^;Trp53^−/−^;Trp53bp1^−/−^* mouse embryonic stem cells (ES-B1P.R mESCs) (Figure S1A). The second screen was performed in *Brca1^−/−^;Trp53^−/−^;Trp53bp1^−/−^* tumor organoids (ORG-KB1P.R), derived from a *K14cre;Brca1^F/F^;Trp53^F/F^* (KB1P) mouse mammary tumor that acquired resistance to PARPi *in vivo* due to loss of 53BP1 function (Duarte *et al*., 2018) (Figure S1A). Both cellular models were transduced with a lentiviral library of 1,976 short hairpin RNA (shRNA) constructs targeting 391 DNA damage response (DDR) related genes (Xu *et al*., 2015; Gogola *et al*., 2018). Cells were either mock treated or selected for 3 weeks in the presence of the PARPi olaparib (Figure 1A). Olaparib selection was carried out at 25nM in ES-B1P.R mESCs and 50nM in ORG-KB1P.R organoids, concentrations which do not affect the viability of resistant cells, but are lethal to the corresponding PARPi-sensitive cells. Sequencing of the shRNAs in the surviving cells revealed a specific and reproducible dropout of hairpins targeting *Lig3* in the olaparib-treated cell population in both ES-B1P.R mESCs and ORG-KB1P.R organoids (Figure1B and S1B, Table S1). Furthermore, *Lig3* was observed to be the only common significant dropout gene identified across both screens (Figure 1C). We therefore decided to investigate further whether LIG3 would constitute a useful target for the reversion of PARPi resistance in BRCA1- deficient cells.

**Figure 1.**
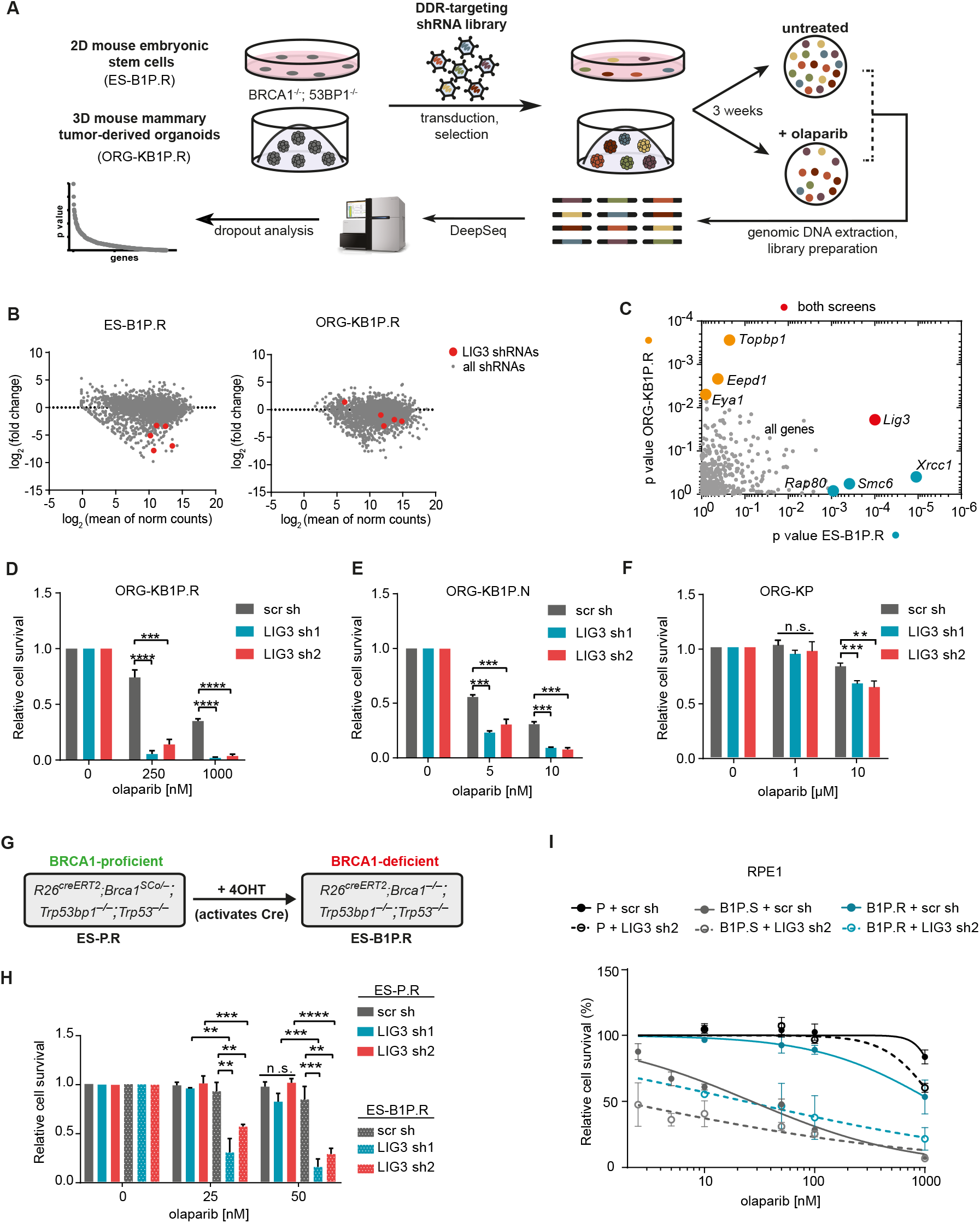
Depletion of LIG3 Increases Sensitivity to PARPi in BRCA1-Deficient Cells, Independent of 53BP1 Loss. **(A)** Outline of the functional shRNA-based dropout screens. The screens were carried out at an olaparib concentration of 25nM and 50nM for ES-B1P.R and ORG-KB1P.R, respectively. **(B)** Plot of Log_2_ ratio (fold change (treated versus untreated)) versus abundance (mean of normalized (norm) counts) of the shRNAs extracted from the screen in ES-B1P.R mESCs and ORG-KB1P.R organoids treated with olaparib or left untreated for three weeks. To eliminate artifacts of significant cell death without PARPi, the analysis of the screens considered fold change between untreated and treated conditions and removed genes that were already depleted in T0 (day of seeding). Analyzed by MAGeCK. **(C)** Comparison of the screening outcome between indicated cell lines, p-value by MAGeCK. **(D-F)** Quantification of long-term clonogenic assays with ORG-KB1P.R **(D)**, ORG-KB1P.S **(E)**, and ORG-KP organoids **(F)** treated with olaparib or left untreated. **(G)** Schematic representation of the *Brca1* selectable conditional allele in *R26^creERT2^;Brca1^SCo/–^;Trp53bp1^−/−^;Trp53^−/−^* (ES-P.R). Incubation of these cells with 4-hydroxytamoxifen (4OHT) induces a CreERT2 recombinase fusion protein, resulting in *R26^creERT2^;Brca1^−/−^;Trp53bp1^−/−^;Trp53^−/−^* (ES- B1P.R) cells lacking BRCA1 protein expression. **(H)** Quantification of long-term clonogenic assay in ES-P.R and ES-B1P.R cells treated with olaparib. See also Figure S2. **(I)** Quantification of long-term clonogenic assays in RPE1-P, RPE1-B1P.S and RPE1-B1P.R cells treated with olaparib. See also Figure S2. Data are represented as mean ± SD. **p<0.01, ***p<0.001, ****p<0.0001, n.s., not significant; two- tailed t test.

### Depletion of LIG3 Increases the Sensitivity to PARPi, Independent of 53BP1 Loss

To validate the findings of our shRNA screens, we carried out viability assays using shRNA- mediated depletion of LIG3 in ORG-KB1P.R organoids. LIG3 depletion significantly increased the sensitivity to olaparib when compared to the parental cells (Figure 1D and S1C). Increased sensitivity to olaparib was also observed upon depletion of LIG3 in PARPi-resistant KB1P.R cells, derived from an independent PARPi-resistant KB1P tumor with 53BP1 loss (Jaspers et al. 2013) (Figure S1A,D,E). These results confirm that depletion of LIG3 results in re-sensitization of BRCA1/53BP1 double-deficient cells to PARPi. Furthermore, depletion of LIG3 also reverted the resistance to olaparib in *Brca1^−/−^;Trp53^−/−^* KB1P.S mammary tumor cells depleted of REV7, a downstream partner of 53BP1 (Boersma *et al*., 2015; Xu *et al*., 2015) (Figure S1A,F,G), indicating that LIG3-mediated resistance is not exclusive for 53BP1-deficient cells.

We next asked whether LIG3 depletion would also increase the PARPi sensitivity of treatment-naïve BRCA1-defcient tumor cells with functional 53BP1. To test this, we used *Brca1^−/−^ ;Trp53^−/−^* organoids, from here onwards refer to as ORG-KB1P.S, and KB1P.S cells derived from independent PARPi-naïve KB1P tumors (Figure S1A) (Jaspers *et al*., 2013; Duarte *et al*., 2018). In both cellular models, shRNA-mediated depletion of LIG3 resulted in increased sensitivity to olaparib (Figure 1E and S1C,H,I). Corroborating our findings, depletion of LIG3 also resulted in increased sensitivity to olaparib in the human *BRCA1*-mutant breast cancer cell line SUM149PT (Figure S1J,K). Importantly, our results were not restricted to olaparib, as LIG3 depletion also increased the sensitivity of KB1P.S cells to the PARPi talazoparib and veliparib (Figure S1L).

### PARPi Sensitization of Cells by LIG3 Depletion is Dependent on BRCA1 Status

Next, we sought to investigate whether the increased PARPi sensitivity of LIG3-depleted cells is BRCA1-dependent. shRNA-mediated depletion of LIG3 in *Trp53^−/−^* organoids (ORG-KP), derived from *K14cre*;*Trp53^F/F^* (KP) mouse mammary tumors (Figure S1A) (Duarte *et al*., 2018), slightly increased the sensitivity to PARPi, but only at a high concentration of 10μM (Figure 1F and S2A,B). To corroborate these data, we validated the effect of LIG3 depletion in *R26^creERT2^;Brca1^SCo/–^;Trp53^−/−^ ;Trp53bp1^−/−^* mESCs (ES-P.R). Addition of 4-hydroxytamoxifen (4OHT) to ES-P.R mESCs induces cre-mediated deletion of the remaining *Brca1* allele, resulting in *R26^creERT2^;Brca1^−/−^;Trp53^−/−^ ;Trp53bp1^−/−^* mESCs (ES-B1P.R), deficient for BRCA1 (Figure 1G and S1A) (Bouwman et al. 2010). Since these mESCs are deficient for p53 and 53BP1, no difference in olaparib sensitivity was observed between the BRCA1-proficient ES-P.R and the BRCA1-deficient ES-B1P.R mESCs (Figure 1H and S2C,D,E,F). shRNA-mediated depletion of LIG3 did not affect cell proliferation in untreated ES-P or ES-B1P.R mESCs. However, LIG3 depletion did result in increased olaparib sensitivity in ES-B1P.R cells, compared to unmodified cells (Figure1H and S2E,F). To investigate whether the effect was independent of the loss of 53BP1, we repeated this experiment in 53BP1- proficient *R26^creERT2^;Brca1^SCo/–^;Trp53^−/−^* mESCs (ES-P) (Figure S1A and S2C,D,G). Depletion of LIG3 increased the sensitivity to PARPi in BRCA1-deficient ES-B1P.S cells but not in BRCA1- proficient ES-P cells (Figure S2H-J).

Additionally, we tested depletion of LIG3 in three isogenic human TERT-immortalized retinal pigment epithelial (RPE1) cell lines with engineered loss of *TP53* (RPE1-P), *TP53*+*BRCA1* (RPE1- B1P.S), or *TP53+BRCA1*+*TP53BP1* (RPE1-B1P.R) (Figure S1A). In line with the data observed in mouse cells, shRNA-mediated depletion of LIG3 only increased sensitivity to olaparib in RPE1-P cells at a higher concentration of 1μM, but rendered RPE1-B1P.R cells as sensitive to olaparib as the RPE1-B1P.S cells (Figure1I and S2K,L). In addition, depletion of LIG3 further increased sensitivity of RPE1-B1P.S cells to olaparib (Figure1I and S2K,L).

Finally, we asked if loss of LIG3 also results in hypersensitization of BRCA2-deficient cells to PARPi. To test this, we used *Brca2^−/−^;Trp53^−/−^* (KB2P) cells derived from a *K14cre;Brca2^F/F^;Trp53^F/F^* (KB2P) mouse mammary tumor (Evers *et al*., 2008) (Figure S1A). shRNA-mediated depletion of LIG3 in KB2P cells resulted in an increase in olaparib sensitivity that was modest compared to the profound increase observed in KB1P cells (Figure S2M,N). In addition, we depleted LIG3 in BRCA2- proficient human DLD1 cells and an isogenic derivative in which *BRCA2* was deleted (DLD1-B2KO). We did not observe a significant increase in sensitivity to olaparib in the BRCA2-deficient DLD1- B2KO cells after depletion of LIG3 (Figure S2O,P). In line with the previous data, depletion of LIG3 in DLD1 cells only resulted in increased olaparib sensitivity at a high concentration of 2.5μM (Figure S2O,Q). Taken together, our data show that LIG3 is a strong modulator of PARPi response specifically in BRCA1-deficient cells and that LIG3 depletion enhances the toxicity of PARPi in BRCA1-deficient cells which acquired resistance due to loss of DSB end-protection.

### PARP1 Trapping Contributes to PARPi Toxicity in LIG3-Depleted Cells

Most PARPi, in addition to blocking the catalytic activity of PARP1, also induce toxic PARP1-DNA complexes as a result of their trapping capacity (Murai *et al*., 2012, 2014). To test whether PARPi- mediated PARP1 trapping contributes to PARPi toxicity in LIG3-depleted cells, we generated *Parp1* knockout isogenic derivatives of ES-P.R mESCs and verified loss of PARP1 expression by western blot (Figure S3A). Compared to cells transduced with non-targeting sgRNA (ES-P.R sgNTG) , ES- P.R-*Parp1^−/−^* cells displayed decreased levels of PAR upon PARG inhibition and/or MMS treatment (Gogola *et al*., 2018), confirming functional loss of PARP1 (Figure S3B). We next exposed ES-P.R sgNTG and ES-P.R-*Parp1^−/−^* cells to 4OHT to produce BRCA1-deficient ES-B1P.R sgNTG and ES- B1P.R-*Parp1^−/−^* mESCs, which were tested for olaparib sensitivity with or without LIG3 depletion (Figure S3C-F). shRNA-mediated depletion of LIG3 did not affect viability of ES-B1P.R sgNTG or ES-B1P.R-*Parp1^−/−^* cells (Figure S3E). In line with the notion that PARPi cytotoxicity is mediated by PARP1 trapping (Murai *et al*., 2012; Pettitt *et al*., 2013), elimination of PARP1 resulted in reduced sensitivity of ES-B1P.R cells to olaparib (Figure S3F). Importantly, elimination of PARP1 also reduced olaparib sensitivity in LIG3-depleted ES-B1P.R cells, indicating that the effect of LIG3 depletion on PARPi sensitivity in BRCA1-deficient cells is partially mediated by PARP1 trapping.

### Resistance to PARPi in 53BP1-deficient KB1P Cells is Mediated by Nuclear LIG3

The *LIG3* gene encodes both mitochondrial and nuclear proteins (Lakshmipathy and Campbell, 1999). Importantly, mitochondrial LIG3 is essential for cellular viability as it ensures mtDNA integrity (Puebla-Osorio *et al*., 2006). Consequently, complete deletion of *Lig3* results in cellular death and early embryonic lethality in mice, whereas nuclear LIG3 has been shown to be dispensable for cell viability (Simsek *et al*., 2011). We therefore asked whether the increased PARPi sensitivity of LIG3- depleted BRCA1-deficient cells resulted from loss of LIG3 activity in the nucleus or in the mitochondria. To test this, we generated nuclear *Lig3* knockout cells which only express the mitochondrial form of LIG3. To this end, we used 53BP1-deficient KB1P.R mouse tumor cells in which we introduced an ATG>CTC mutation in the internal translation initiation site that is required for expression of the nuclear LIG3 isoform but does not affect expression of mitochondrial LIG3 (Figure 2A) (Lakshmipathy and Campbell, 1999). Western blot analysis of KB1P.R cells, one KB1P.R(LIG3^mut/wt^) clone heterozygous for the ATG>CTC mutation (B1) and two KB1P.R(LIG3^mut/mut^) clones with homozygous ATG>CTC mutation (A3, F5) showed that LIG3 is still expressed (Figure 2B). However, immunofluorescence analysis of LIG3 in the same clones revealed that parental KB1P.R cells and the heterozygous KB1P.R(LIG3^mut/wt^) B1 clone displayed LIG3 staining in both nucleus and mitochondria, whereas the homozygous KB1P.R(LIG3^mut/mut^) A3 and KB1P.R(LIG3^mut/mut^) F5 clones exhibited loss of nuclear LIG3 expression (Figure 2C). Finally, we investigated whether the nuclear mutants of LIG3 displayed increased sensitivity to PARPi. Long-term clonogenic assays revealed that the nuclear LIG3-deficient KB1P.R(LIG3^mut/mut^) A3 and KB1P.R(LIG3^mut/mut^) F5 clones displayed hyper-sensitivity to olaparib when compared to the PARPi-resistant parental KB1P.R cells and the heterozygous KB1P.R(LIG3^mut/wt^) B1 clone (Figure 2D and S4A,B).

**Figure 2.**
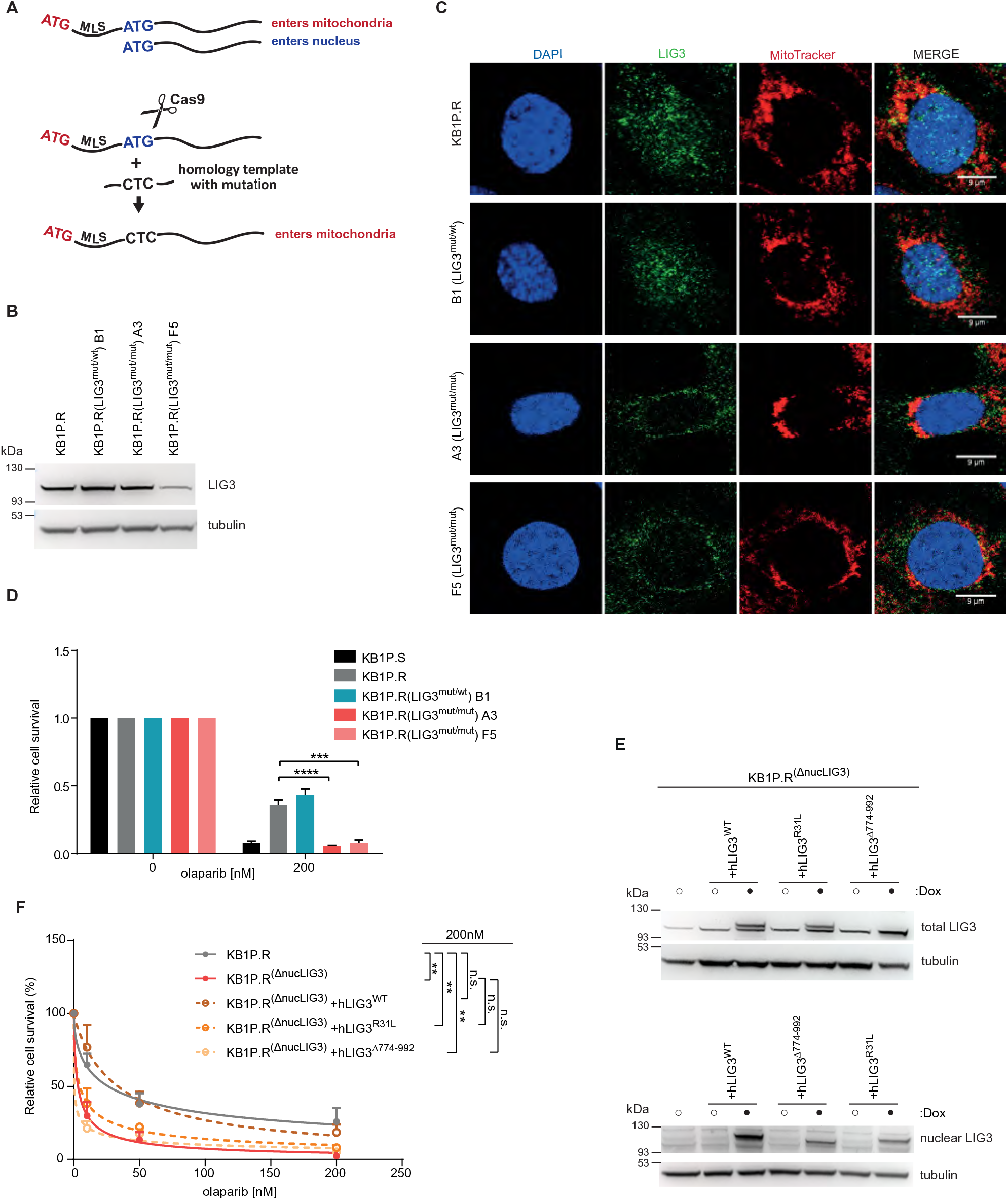
Resistance to PARPi in 53BP1-deficient KB1P cells is Mediated by Nuclear LIG3. **(A)** Schematic representation of the generation of nuclear LIG3 mutants in KB1P.R cells. *Lig3* contains two translation initiation ATG sites. The sequence flanked by both ATG sites functions as a mitochondrial targeting sequence. If translation is initiated at the upstream ATG site, a mitochondrial protein is produced, whereas if translation initiated at the downstream ATG site produces the nuclear form. Ablation of the downstream ATG allows cells to retain mitochondrial LIG3 function, but not nuclear function. CRISPR/Cas9 system was used to introduce in-frame ATG>CTC mutation in the nuclear ATG through the delivery of a homology repair template. **(B)** Western blot analysis of LIG3 in whole cell lysates of KB1P.R, KB1P.R(LIG3^mut/wt^) B1, KB1P.R(LIG3^mut/mut^) A3 and KB1P.R(LIG3^mut/mut^) F5 cells. **(C)** Immunofluorescence of LIG3 together with MitoTracker staining to examine the sub-cellular localization of LIG3 in mutant cells. **(D)** Quantification of long-term clonogenic assays with KB1P.S, KB1P.R cells and nuclear LIG3 mutant clones B1, A3 and F5, treated with olaparib or untreated. **(E)** Western blot analysis of total and nuclear LIG3 in KB1P.R and nuclear LIG3 mutant KB1P.R^(ΔnucLIG3)^ cells. Expression of LIG3 constructs was induced with Doxycycline (Dox) for 2 days prior to analysis. **(F)** Quantification of long-term clonogenic assay with KB1P.R and nuclear LIG3 mutant KB1P.R^(ΔnucLIG3)^ cells, treated with olaparib or untreated. Expression of LIG3 constructs was induced with Doxycycline (Dox) starting 2 days before the assay and maintained for the duration of the assay. Data are represented as mean ± SD. **p<0.01, ***p<0.001, ****p<0.0001; two-tailed t test.

Nuclear LIG3 consists of a N-terminal like zinc finger (ZnF) domain which is required for binding to DNA secondary structures (Taylor, Whitehouse and Caldecott, 2000) and a C-terminal BRCT domain required for interaction with other proteins such as XRCC1 (Caldecott *et al*., 1994). To test the role of these domains in LIG3-mediated PARPi resistance, we generated overexpression constructs for wild-type human LIG3 (hLIG3^WT^), carrying a mutation in the PARP-like ZnF domain (hLIG3^R31L^) or a C-terminal Δ774-922 truncation (hLIG3^Δ774-922^). We introduced these constructs in KB1P.R(LIG3^mut/mut^) A3 cells - from here onwards referred to as KB1P.R^(ΔnucLIG3)^ - and carried out clonogenic assays (Figure 2E and S4C). Whereas overexpression of hLIG3^WT^ rescued sensitivity to olaparib in KB1P.R^(ΔnucLIG3)^ cells, overexpression of either hLIG3 mutant failed to suppress olaparib sensitivity in KB1P.R^(ΔnucLIG3)^ cells (Figure 2F AND S4C), indicating that both the DNA binding and BRCT domain are required for driving PARPi resistance in BRCA1 and 53BP1 double-deficient tumor cells.

### LIG3 is Required at Replication Forks in BRCA1-Deficient Cells Treated with PARPi

Our data indicates that the increase in sensitivity to PARPi arising from LIG3 depletion is independent of the loss of DSB end-protection and therefore we hypothesized that this phenomenon could be independent of HR status. To test this hypothesis, we carried out RAD51 ionizing radiation-induced foci (RAD51 IRIF) in our mouse tumor-derived cell lines as a read-out of functional HR status (Xu *et al*., 2015). As expected, BRCA1-deficient KB1P.S cells had significantly less IRIF per cell than the BRCA1-proficient KP cells (Figure S1A), while the BRCA1/53BP1 double- deficent KB1P.R cells displayed increased numbers of IRIF compared with KB1P.S (Figure S5A). Moreover, KB1P.R cells with shRNA-mediated depletion of LIG3 or with deletion of LIG3 nuclear isoform did not show a significant reduction of RAD51 IRIF (Figure S5A), corroborating our hypothesis that the sensitivity observed in LIG3-depleted cells is not a result of decreased HR in these cells.

LIG3 is also involved in the repair of DSBs by alternative end-joining (Alt-EJ) through its interaction with POLθ (Wang *et al*., 2005; Simsek *et al*., 2011). It has been previously reported that HR-deficient tumors rely on POLθ for survival and that its depletion can enhance PARPi-response in both BRCA1-deficient cells and BRCA1/53BP1 double-mutant cells (Ceccaldi *et al*., 2015; Mateos-Gomez *et al*., 2015; Zhou *et al*., 2020). Therefore, we hypothesized that, if the suppressive effect of LIG3 on PARPi sensitivity in BRCA1-deficient cells is dependent on its role in Alt-EJ, viability of LIG3-deficient cells should not be affected by inhibition of POLθ and that sensitivity of LIG3-deficient cells to olaparib would not be amplified by POLθ inhibition. To test this, we carried out both long-term clonogenic and short-term cytotoxicity assays with olaparib and the POLθ inhibitor ART558 (Zatreanu *et al*., 2021). Interestingly, inhibition of POLθ alone resulted in increased cell death in KB1P.R parental cells, as well as in nuclear LIG3 mutant KB1P.R^(ΔnucLIG3)^ cells (Figure S5B,C). Moreover, we observed a synergistic interaction between olaparib and ART558 in both cell lines (Figure S5C), suggesting that LIG3-mediated resistance is independent of its role in POLθ- mediated end-joining.

Data from recent studies indicate that LIG3 is present at replication forks (Arakawa and Iliakis, 2015; Hanzlikova *et al*., 2018; Sriramachandran *et al*., 2020; Cong *et al*., 2021). Therefore, we next investigated whether LIG3 localizes to sites of DNA replication marked by 5-ethynyl-2’- deoxyuridine (EdU) incorporation, in the absence of DNA damage induction. To test this, we performed proximity ligation-based assays (PLA) to detect LIG3 binding to replicated DNA (Taglialatela *et al*., 2017; Mukherjee *et al*., 2019), in BRCA1-proficient KP, BRCA1-reconstituted KB1P.S+hB1 (Barazas *et al*., 2019), and in BRCA1-deficient KB1P.S cells. Interestingly, untreated KB1P.S cells showed significantly higher levels of LIG3-EdU PLA foci than KP or KB1P.S+hB1cells (Figure 3A,B and S5D,E). We next tested if LIG3 localization at replication sites is affected by PARPi treatments which would trap PARP1 at RFs. Therefore, we carried out LIG3-EdU PLA after incubating cells with olaparib for 2hr. Quantification of LIG3-EdU PLA foci revealed that PARPi treatment did not induce any increase in the number of foci in KB1P.S+hB1 cells. In contrast, BRCA1-deficient KB1P.S cells displayed a striking increase in the number of PLA foci after olaparib treatment (Figure 3A and S5D). We next investigated whether LIG3 localization at replication sites is affected by the PARG inhibitor (PARGi) PDDX-001 which is known to increase PAR levels and to also result in an increase in chromatin-associated PARP1 (James *et al*., 2016; Gogola *et al*., 2018; Hanzlikova *et al*., 2018). We therefore carried out LIG3-EdU PLA after incubating cells with PDDX- 001 for 30 min. Similar to olaparib-treated cells, PDDX-001-treated BRCA1-deficient cells showed a strong increase in the number of LIG3-EdU PLA foci, while no significant changes were observed in KP cells (Figure 3B and S5E). Co-localization of LIG3 at EdU-marked replication sites after PDDX-001 treatment was also verified qualitatively by LIG3 immunostaining in KP, KB1P.S cells and KB1P.R cells (Figure S5F). Since we observe that both PARPi and PARGi treatment results in an increase in LIG3-EdU PLA foci, and that olaparib treatment results in a reduction of PAR levels while treatment with PDDX-001 results in an increase, we conclude that the upsurge in LIG3-EdU PLA is probably caused by PARP1 trapping, which is common to both inhibitors. Of note, untreated KB2P cells showed similar numbers of LIG3-EdU PLA foci as KP cells (Figure S5G,H). However, upon treatment with PDDX-001, KB2P cells showed more LIG3-EdU PLA foci than KP cells, but significantly less than KB1P.S cells (Figure S5G,H). These data support our previous findings that LIG3 depletion in BRCA2-deficient cells has a more modest effect on olaparib sensitivity than in BRCA1-deficient cells.

**Figure 3.**
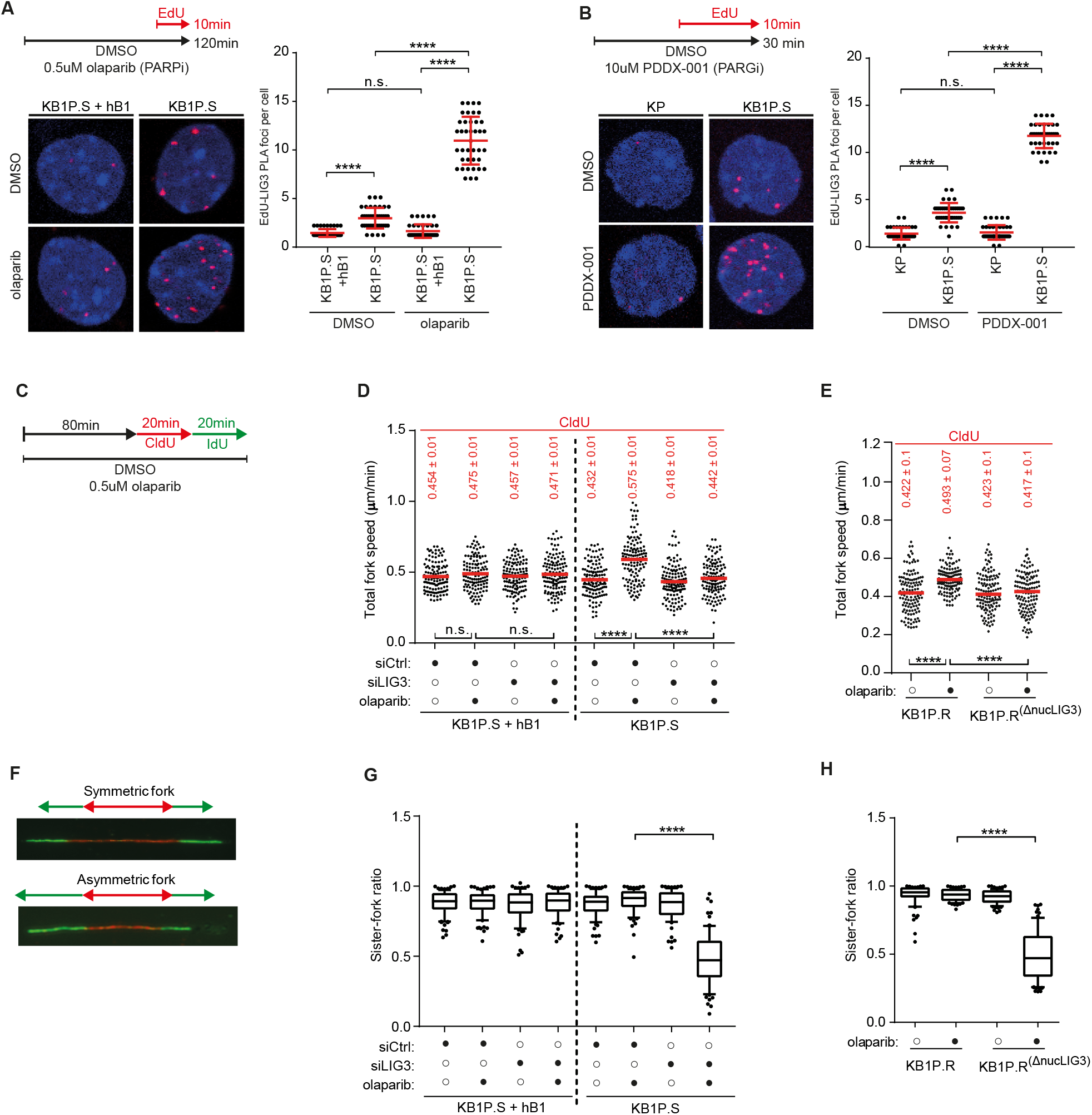
LIG3 is Required at Replication Forks in BRCA1-Deficient Cells Treated with PARPi. **(A)** Outline of experimental set up, representative images and quantification of LIG3-EdU proximity ligation assay (PLA) foci, in KB1P.S+hB1 and KB1P.S cells incubated for 10 min with 20μM EdU, in the absence or presence of 0.5μM olaparib. **(B)** Outline of experimental set up, representative images and quantification of LIG3-EdU PLA foci in KP and KB1P.S cells incubated for 10 min with 20μM EdU, in the absence or presence of PDDX- 001. **(C)** Outline of DNA fiber assay experimental set up and representative images of DNA replication forks. Cells were pre-incubated with 0.5μM olaparib for 80 min, followed by sequential labeling with CldU (red) and IdU (green) in the presence of olaparib for 20 min each. Replication fork progression was quantified by measuring tract lengths of CldU and IdU in micrometers (μM). **(D)** Quantification of fork speed in CldU tracks, following the indicated treatments, in KB1P.S+hB1 and KB1P.S cells after siRNA-mediated depletion of LIG3. See also Figure S6. **(E)** Quantification of fork speed in CldU tracks, following the indicated treatments, in nuclear LIG3 mutant KB1P.R^(ΔnucLIG3)^ cells. See also Figure S6. **(F)** Representative images of symmetric and asymmetric replication forks. **(G)** Quantification of fork symmetry following the indicated treatments in KB1P.R cells. The box represents the 10th to 90th percentiles. **(H)** Quantification of fork symmetry following the indicated treatments in KB1P.R and KB1P.R^(ΔnucLIG3)^ cells. The box represents the 10th to 90th percentiles. Data are represented as mean. ****p<0.0001; n.s., not significant; Mann–Whitney U test.

Since LIG3 seems to play a role at replication sites in BRCA1-deficient conditions, we asked whether depletion of LIG3 would affect RF progression in untreated and PARPi-treated BRCA1- deficient cells. To test this, we performed DNA fiber assay in BRCA1-deficient KB1P.S and BRCA1- reconstituted KB1P.S+hB1 cells. Cells were pre-incubated with olaparib for 80 min, followed by sequential labelling with CldU (red) and IdU (green) for 20 mins each in the presence of olaparib (Figure 3C). Progression was measured by tract lengths of CldU and IdU. Analysis of RF speeds revealed no significant increase in BRCA1-proficient KB1P.S+hB1 cells after olaparib treatment (Figure 3D and S5I,J). In contrast, BRCA1-deficient KB1P.S cells exhibited an increase in RF speed upon olaparib treatment, in line with previous work (Cong *et al*., 2021). Surprisingly, while siRNA- mediated depletion of LIG3 did not affect RF speed in untreated cells, it significantly suppressed the PARPi-induced increase in fork speed in KB1P.S cells (Figure 3D and S5I,J). As observed in KB1P.S cells, olaparib treatment also resulted in increased RF speed in BRCA1/53BP1 double-deficent KB1P.R cells, which was rescued by siRNA-mediated LIG3 depletion or loss of nuclear LIG3 (Figure 3E and S5I,K-M). Similar data was also observed in BRCA1/53BP1 double-deficient RPE1-B1P.R cells treated with PARPi (Figure S5N,O). Since loss of LIG3 rescued the increase in fork speed in both BRCA1 and BRCA1/53BP1 double-deficient cells, we asked if this phenomenon was due to a restraint in fork speed or due to continuous fork stalling and restart and thus increased replication stress. We next analyzed RF symmetry in BRCA1-proficient and-deficient cells by measuring sister fork-ratio (Figure 3F,G). While BRCA1-proficient KB1P.S+hB1 cells did not show any significant differences in fork symmetry across conditions, depletion of LIG3 induced a significant increase in sister fork asymmetry, indicative of fork stalling, in BRCA1-deficient KB1P.S cells exposed to olaparib (Figure 3G). Similarly, loss of nuclear LIG3 in KB1P.R^(ΔnucLIG3)^ cells also resulted in fork asymmetry upon olaparib treatment (Figure 3H). These data corroborate our hypothesis that the lack of PARPi-induced fork acceleration observed in LIG3-depleted cells is a result of persistent RF stress upon loss of LIG3. Overall, our results support the notion that depletion of LIG3 in BRCA1-deficient cells exposed to PARPi leads to slower and asymmetric forks.

### Loss of LIG3 in BRCA1-Deficient Cells Results in an Increase in PARPi-mediated ssDNA Regions

Recent studies have suggested that accumulation of post-replicative single-stranded DNA (ssDNA) gaps underlies BRCA deficiency and PARPi sensitivity (Quinet *et al*., 2020; Cong *et al*., 2021; Panzarino *et al*., 2021). Since LIG3 is a DNA ligase and our data indicates that it is present at active RFs in BRCA1-deficient cells, we asked whether LIG3 depletion would result in an increase in S phase associated ssDNA. To test this, we cultured KB1P.S+hB1, KB1P.S and KB1P.R mouse tumor cells in medium supplemented with BrdU for 48hr followed by a 2hr-treatment with olaparib and quantification of native BrdU intensity by quantitative image-based cytometry (QIBC) (Toledo *et al*., 2013) (Figure 4A). As previously suggested, olaparib treatment did not result in an increase in ssDNA levels in BRCA1-proficient KB1P.S+hB1 cells nor in BRCA1/53BP1 double-deficient KB1P.R cells (Figure 4B,C and S6A,B). However, treatment with olaparib resulted in a significant increase in ssDNA levels in BRCA1-deficient KB1P.S cells during S-phase (Figure 4B,C, and S6A,B). These results were further confirmed in the RPE1 isogenic lines, showing PARPi-induced increase in ssDNA levels in RPE1-B1P.S cells but not in RPE1-P or RPE1-B1P.R cells (Figure S6D). Importantly, deletion of nuclear LIG3 in KB1P.R cells or shRNA-mediated LIG3 depletion in RPE1- B1P.R cells restored PARPi-induced ssDNA gaps accumulation (Figure 4D, and S6C,D). LIG3 depletion also further increased PARPi-induced ssDNA gaps accumulation in RPE1-B1P.S cells (Figure S6D), suggesting that LIG3-mediated ssDNA gap suppression is HR-independent.

**Figure 4.**
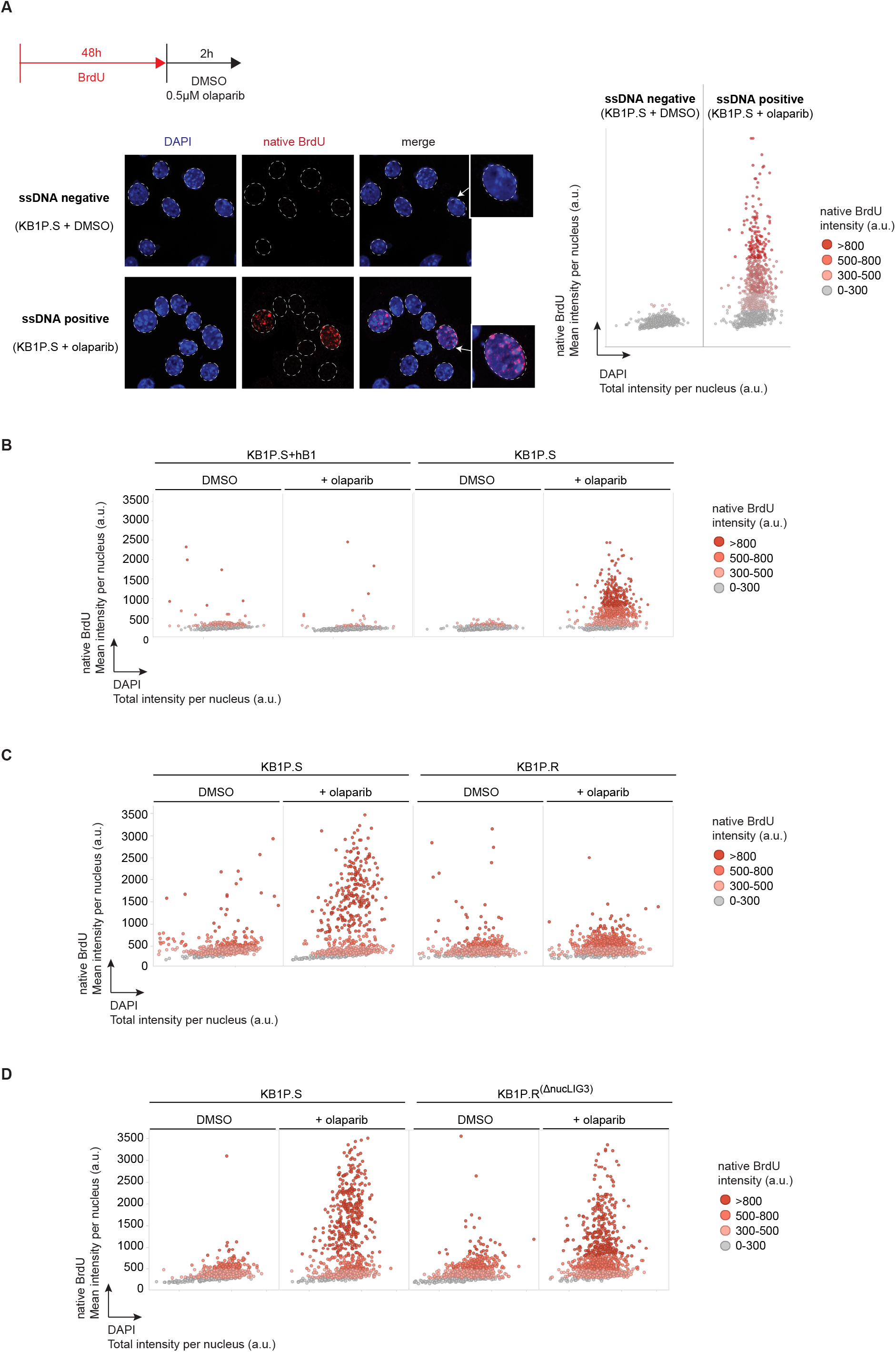
Loss of LIG3 in BRCA1-Deficient Cells Results in an Increase in PARPi-mediated ssDNA Regions. **(A)** Outline of experimental set up to quantify amount of ssDNA gaps per nucleus by quantitative image-based cytometry (QIBC) analysis of mean intensity of native BrdU per nucleus. Cells were incubated with BrdU for 48 hr followed by 2 hr treatment with 0.5μM olaparib or left untreated. **(B)** QIBC analysis of ssDNA in KB1P.S+hB1 and KB1P.S cells. **(C)** QIBC analysis of ssDNA in KB1P.S and KB1P.R cells. **(D)** QIBC analysis of ssDNA in KB1P.S and LIG3 nuclear mutant KB1P.R^(ΔnucLIG3)^ cells. See also Figure S7.

### Increase in ssDNA Gaps Results in Increased Genomic Instability in LIG3-deficient Cells

MRE11 has been shown to be involved in the processing of gaps at and behind DNA replication forks (Hashimoto *et al*., 2010; Schlacher *et al*., 2011; Ray Chaudhuri *et al*., 2016). Furthermore, the nucleosome remodeling factor CHD4 has been reported to be involved in the recruitment of MRE11 for nuclease processing at stressed forks (Ray Chaudhuri *et al*., 2016). We therefore tested if the PARPi-induced increase in replication-associated ssDNA regions in KB1P.S and KB1P.R^(ΔnucLIG3)^ cells was dependent on either MRE11 or CHD4. Both inhibition of MRE11 with mirin and siRNA-mediated depletion of CHD4 rescued the increase in replication-associated ssDNA regions in KB1P.R^(ΔnucLIG3)^ cells treated with olaparib (Figure 5A and S6E-G). In contrast, neither treatment with mirin nor depletion of CHD4 rescued ssDNA exposure in parental KB1P.S cells (Figure 5A and S6F,G). To confirm if the observed increase of ssDNA was in the vicinity of RFs, we used electron microscopy (EM) to visualize the fine architecture of replication intermediates in KB1P.S, KB1P.R and KB1P.R^(ΔnucLIG3)^ cells after 2hr-treatment with olaparib (Figure 5B,C). In untreated conditions, a minority of the DNA molecules displayed ssDNA gaps behind the fork in all the three cell lines analyzed. However, olaparib treatment markedly enhanced the percentage of molecules displaying 1 or more post-replicative ssDNA gaps, specifically in KB1P.S and KB1P.R^(ΔnucLIG3)^ but not in KB1P.R cells (Figure 5D). Consistent with our QIBC data, we observed that the PARPi-induced post-replicative gaps in KB1P.S cells were not rescued upon inhibition of MRE11 whereas the post-replicative gaps in olaparib-treated KB1P.R^(ΔnucLIG3)^ cells were dependent on MRE11-mediated processing (Figure 5D). Of note, we did not observe an increase in fork reversal in any of the conditions (Figure S6H). Taken together, these data suggest that the PARPi-induced ssDNA regions in BRCA1-deficient and 53BP1-proficient cells are distinct in nature from the PARPi-induced gaps generated upon loss of LIG3 in BRCA1/53BP1 double-deficient cells.

**Figure 5.**
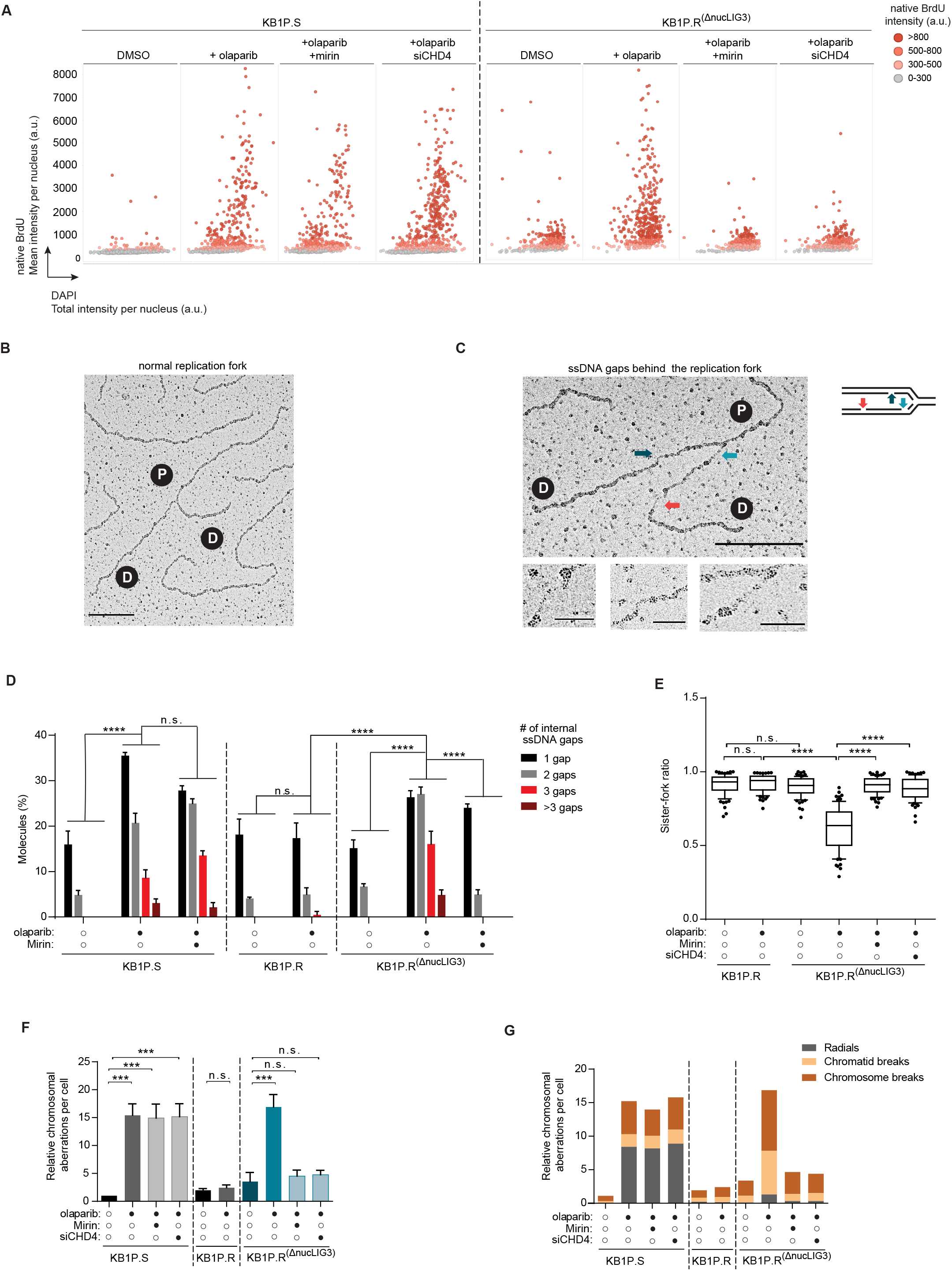
Increase in ssDNA Gaps Results in Increased Genomic Instability in LIG3-deficient cells. **(A)** QIBC analysis of ssDNA gaps in KB1P.S and nuclear LIG3 mutant KB1P.R^(ΔnucLIG3)^ cells. Cells were treated with 25μM mirin for 48hr prior to treatment with olaparib, or transfected with siRNA targeting CHD4. See also Figure S7. **(B and C)** Representative electron micrographs of normal replication fork **(B)** and fork with internal ssDNA gaps behind replication fork **(C)**. Scale bar for large panels: 250nm = 1214bp; scale bar for small panels: 50nm = 242bp. P; parental strand. D; daughter strand. **(D)** Quantification of internal ssDNA gaps behind replication forks observed in KB1P.S, KB1P.R and KB1P.R^(ΔnucLIG3)^ cells upon treatment with 0.5μM olaparib for 2hr. KB1P.S and KB1P.R^(ΔnucLIG3)^ cells were additionally treated with 25μM mirin for 48hr prior to treatment with olaparib, or transfected with siRNA targeting CHD4. Data were acquired by electron microscopy. Data are represented as mean ± SD. ****p<0.0001, n.s., not significant; two-way ANOVA. **(E)** Quantification of fork symmetry in KB1P.R and KB1P.R^(ΔnucLIG3)^ cells following the indicated treatments. KB1P.R^(ΔnucLIG3)^ cells were additionally treated with 25μM mirin for 48hr prior to treatment with olaparib, or transfected with siRNA targeting CHD4. Data are represented as mean and the box represents the 10th to 90th percentiles. ****p<0.0001; n.s., not significant; Mann– Whitney U test. **(F)** Quantification of chromosomal aberrations in KB1P.S, KB1P.R and KB1P.R^(ΔnucLIG3)^ cells following 2 hr treatment with 0.5μM olaparib and recovery for 6 hr. KB1P.S and KB1P.R^(ΔnucLIG3)^ cells were additionally treated with 25μM mirin for 48hr prior to treatment with olaparib, or transfected with siRNA targeting CHD4. Data are represented as mean ± SD. ***p<0.001, n.s., not significant; two-tailed t test. **(G)** Quantification of the different types of chromosomal aberrations identified in (F).

We next questioned if the suppression of post-replicative gaps observed upon either MRE11 inhibition or depletion of CHD4 could result in fork stability in KB1P.R^(ΔnucLIG3)^ cells. To assess this, we performed DNA fiber assays to measure fork asymmetry in KB1P.R and KB1P.R^(ΔnucLIG3)^ cells upon exposure to olaparib combined with either MRE11 inhibition or CHD4 depletion. Interestingly, our data revealed that the fork asymmetry observed in these cells upon treatments with olaparib was completely rescued upon MRE11 inhibition or depletion of CHD4 (Figure 5E). However, MRE11 inhibition or CHD4 depletion did not result in an increase in fork speed in KB1P.R^(ΔnucLIG3)^ cells exposed to olaparib as observed in KB1P.R cells, suggesting that the increase in fork speed is uncoupled from MRE11-mediated ssDNA gap exposure and from PARPi sensitivity (Figure S6I,J).

We next tested whether the increase in post-replicative ssDNA gaps upon LIG3 depletion resulted in increased genomic instability. We analyzed chromosomal aberrations in metaphase spreads of KB1P.S+hB1, KB1P.S, KB1P.R and KB1P.R^(ΔnucLIG3)^ cells after treatment with olaparib for 2hr. As expected, olaparib treatment resulted in increased numbers of chromosomal aberrations in KB1P.S cells but not in KB1P.S+hB1 and KB1P.R (Bunting *et al*., 2010) (Figure 5F and S7K). Interestingly, KB1P.R^(ΔnucLIG3)^ cells showed a surge in chromosomal aberrations when compared to KB1P.R cells (Figure 5F,G). Interestingly, the aberrations in PARPi-treated KB1P.R^(ΔnucLIG3)^ cells mainly consisted of chromosome and chromatid breaks, whereas PARPi-treated KB1P.S cells showed more radials (Figure 5G). siRNA-mediated depletion of LIG3 further enhanced chromosomal aberrations in KB1P.S cells (Figure S6K). Of note, inhibition of MRE11 with mirin or siRNA-mediated depletion of CHD4 suppressed PARPi-induced genomic instability in KB1P.R^(ΔnucLIG3)^ cells, indicating that PARPi-induced genomic instability in these cells is mediated by MRE11-dependent ssDNA gap exposure (Figure 5F,G and S6L). As expected, treatment with mirin or depletion of CHD4 did not rescued chromosomal aberrations in parental KB1P.S cells (Figure 5F,G). Importantly, loss of LIG3 did not result in an increase in immediate DSBs following olaparib treatment, as assessed by pulsed-field gel electrophoresis (PFGE) of genomic DNA from KB1P.S+hB1, KB1P.S and KB1P.R cells and by immunofluorescence analysis of γ-H2AX foci in K.P, KB1P.S and KB1P.R cells (Figure S7A,B). Altogether, these data indicate that the increase in genomic instability induced by loss of nuclear LIG3 in BRCA1/53BP1 double-deficient cells exposed to PARPi is caused by post-replicative ssDNA gaps.

### LIG3 Depletion Increases *in vivo* Efficacy of PARPi

Our previous results established that LIG3 is a modulator of PARPi-response *in vitro*. To test whether our results could be recapitulated *in vivo*, we performed shRNA-mediated depletion of LIG3 in PARPi-naïve KB1P4.N1 organoids (BRCA1-deficient) and PARPi-resistant KB1P4.R1 organoids (BRCA1/53BP1 double-deficient) (Figure 6A and S1A). The modified organoid lines were transplanted into the mammary fat pad of syngeneic wild-type mice. Upon tumor outgrowth, mice were treated with olaparib or vehicle for 28 consecutive days, and mice were sacrificed when tumors progressed to a volume of ≥1500 mm^3^. LIG3 depletion did not affect tumor growth and all cohorts of vehicle-treated mice showed comparable survival (Figure 6B,C). In contrast, LIG3 depletion significantly enhanced the anticancer efficacy of olaparib, resulting in increased survival of olaparib-treated mice bearing KB1P4.N1+shLIG3 tumors, compared to olaparib-treated mice with KB1P4.N1+shscr tumors (Figure 6B). Importantly, LIG3 depletion also resensitized the PARPi-resistant KB1P4.R1 tumors to olaparib. Whereas olaparib-treated and vehicle-treated mice with KB1P4.R1 tumors showed comparable survival, olaparib treatment significantly prolonged the survival of mice bearing KB1P4.R1+shLIG3 tumors (Figure 6C). Together, these data show that LIG3 also modulates PARPi response *in vivo*.

**Figure 6.**
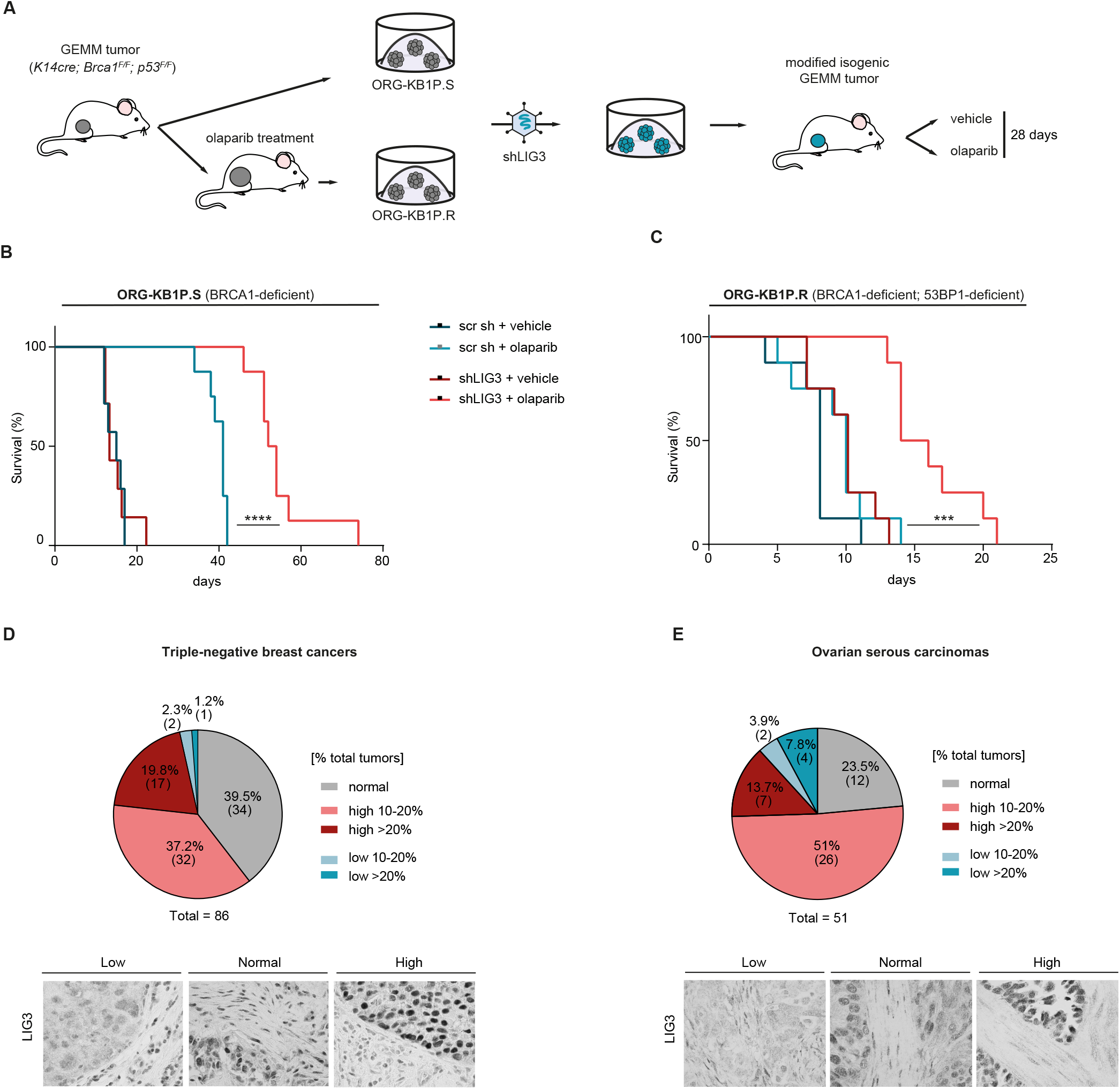
LIG3 Depletion Increases *in vivo* Efficacy of PARPi and is Overexpressed in a Fraction of Human Tumors. **(A)** Schematic outline of *in vivo* experimental set up. Organoids were modified *in vitro* and transplanted into the mammary fat pad of syngeneic, wild-type FVB/NRj mice. Upon tumor outgrowth, mice were treated with olaparib or vehicle for 28 consecutive days. **(B and C)** Kaplan–Meier survival curves of mice transplanted with KB1P.S **(B)** or KB1P.R organoid lines **(C)**, after *in vitro* shRNA-mediated depletion of LIG3. ***p<0.001, ****p<0.0001; Log-Rank (Mantel Cox). **(D and E)** Summary and representative images of immunohistochemistry (IHC) analysis of LIG3 expression in triple-negative breast cancers **(D)** and ovarian serous carcinomas **(E)**.

### Increased LIG3 Expression in Triple-Negative Breast and Serous Ovarian Cancers

To assess the clinical relevance of LIG3, we determined LIG3 expression in sections of treatment-naïve tumors from a cohort of 86 women with triple-negative breast cancer (TNBC) (Gogola *et al*., 2018) and 51 women with high-grade serous ovarian carcinoma (Moudry *et al*., 2016), two clinically relevant groups of patient eligible for PARPi treatment. Immunohistochemistry (IHC) analysis revealed that, while LIG3 protein was expressed at normal levels in a majority of tumor cells in the biopsies, a substantial proportion of samples contained areas displaying aberrant expression of LIG3. Of the 87 TNBC cases analyzed, 32 (37.2%) and 17(19.8%) biopsies showed LIG3 overexpression in areas corresponding to >10% and >20% of the tumor, respectively (Figure 6D). Similarly, 26 (51%) and 7 (13.7%) of the 51 ovarian cancer cases showed LIG3 overexpression in areas corresponding to >10% and >20% of the tumor, respectively (Figure 6E). Conversely, LIG3- negative areas were observed in a small proportion of biopsies, with 2 (2.3%) and 1 (1.2%) of the 86 TNBC cases, and 2 (3.9%) and 4 (7.8%) of the 51 ovarian cancers displaying loss of LIG3 in areas corresponding to >10% and >20% of the tumor, respectively (Figure 6D,E). These observations reveal that LIG3 expression is heterogeneous within and across TNBC and serous ovarian cancers, which might result in selective expansion of LIG3 overexpressing clones during PARPi treatment and thereby contribute to intratumoral and inter-patient differences in response to PARPi therapy.

## DISCUSSION

Molecular alterations that render cells resistant to targeted therapies may also cause synthetic dependencies, which can be exploited to design rational combination therapies. However, the pathways that can be targeted to exploit these vulnerabilities are poorly understood. In this study, we used shRNA dropout screens to identify synthetic dependencies of BRCA1-deficient cells which acquired resistance to PARPi treatment by restoration of HR due to loss 53BP1. We have identified LIG3 as a critical suppressor of PARPi toxicity in BRCA1/53BP1 double-deficient cells. Loss of LIG3 also enhances PARPi sensitivity of BRCA1-deficient cells with intact 53BP1, indicating that the role of LIG3 in BRCA1-deficient cells is independent of their 53BP1 status.

In this study, we show that the increase in sensitivity to PARPi observed upon LIG3 loss in BRCA1/53BP1 double-deficient cells results from an increase in post-replicative MRE11-dependent ssDNA gaps. This is in line with the notion that PARPi treatment results in accumulation of post-replicative ssDNA gaps and that exposure to these lesions is a key determinant of PARPi response (Quinet *et al*., 2020; Cong *et al*., 2021). Moreover, our data show that exposure to post-replicative ssDNA gaps underlies PARPi cytotoxicity in both HR-deficient and HR-restored cells, indicating that LIG3-mediated PARPi resistance in BRCA1/53BP1 double-deficient cells is an HR-independent mechanism. Together, these data indicate that BRCA1/53BP1 double-deficient cells rely on LIG3 for suppression of PARPi-induced gaps, rendering LIG3 as a synthetic dependency of these cells. LIG3 depletion also increased sensitivity to PARPi in BRCA1/REV7 double-deficient cells, suggesting this synthetic dependency is common to BRCA1-deficient tumor cells that acquired PARPi resistance due to loss of end-protection.

We show that PARPi-induced ssDNA gaps in BRCA1-deficient cells are not substrates for MRE11-mediated degradation, indicating that PARPi-induced ssDNA gaps observed in LIG3- depleted BRCA1/53BP1 double-deficient cells are distinct from the gaps in BRCA1-deficient cells. Together, these data suggest existence of two different mechanisms of gap suppression in BRCA1- deficient cells, one dependent on loss of 53BP1 and another which is LIG3-dependent. In PARPi-sensitive BRCA1-deficient cells, 53BP1 drives the formation of post replicative ssDNA gaps upon PARPi treatment. Loss of LIG3 in these cells further enhances accumulation of PARPi-induced ssDNA gaps. On the other hand, PARPi-resistant BRCA1/53BP1 double-deficient cells are competent for HR and thus lack 53BP1-mediated gap formation, hence PARPi-induced ssDNA gaps only occur upon loss of LIG3 (Figure 7).

**Figure 7.**
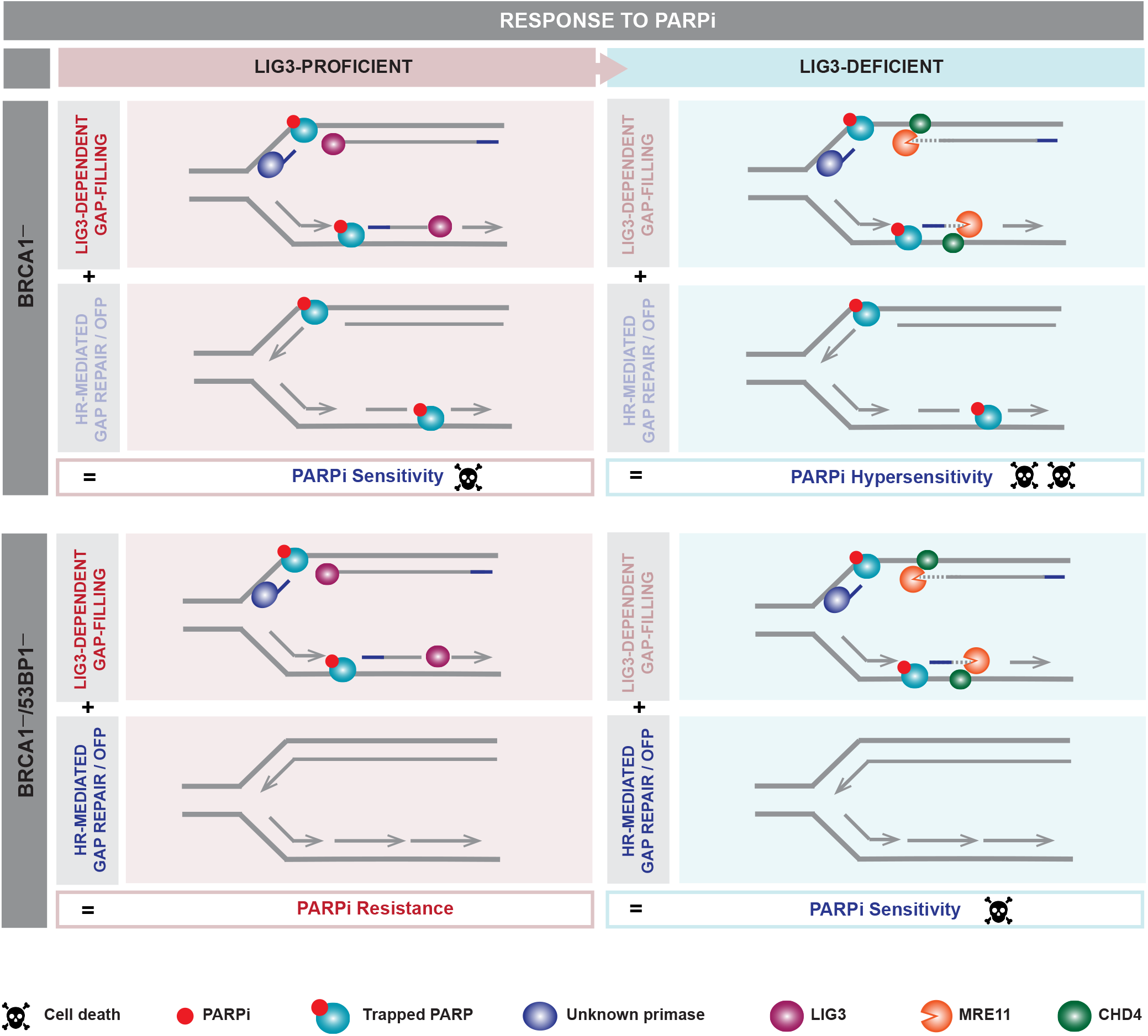
Proposed Model. In response to chromatin-trapped PARP1 lesions, BRCA1-deficient cells have two different mechanisms of gap suppression required for lesion bypass: one dependent on loss of 53BP1 and another which is LIG3-dependent. 53BP1-mediated ssDNA gap induction may result from loss of homologous recombination (HR)-mediated gap repair and/or defective Okazaki fragment processing. LIG3-mediated gap suppression might require repriming activities mediated by Polα, PRIMPOL or another unknown primase, resulting in small gaps which depend on LIG3 to be filled. Upon loss of LIG3, recruitment of MRE11 by CHD4 leads to unscheduled processing of the small gaps into longer stretches of post-replicative ssDNA, resulting in fork stalling and increased genomic instability. PARPi-sensitive BRCA1-deficient cells exhibit post-replicative PARPi-induced ssDNA gaps which are mediated by 53BP1. Accumulation of PARPi-induced post-replicative ssDNA gaps mediated by 53BP1 and by loss of LIG3 underlies PARPi hypersensitivity of BRCA1/LIG3 double-deficient cells. Conversely, PARPi-resistant BRCA1/53BP1 double-deficient cells lack 53BP1-mediated gap formation, and PARPi-induced ssDNA gaps only occur upon loss of LIG3, resulting in accumulation of longer stretches of post-replicative ssDNA, ultimately leading to fork stalling, genomic instability and rendering cells sensitive to PARPi.

53BP1-mediated gap induction in BRCA1-deficient cells exposed to PARPi may result from loss of recombinatorial gap repair (Branzei and Szakal, 2016) and/or defective Okazaki fragment processing (OFP) due to loss of the PARP1-XRCC1-LIG3 backup pathway (Arakawa and Iliakis, 2015; Hanzlikova *et al*., 2018; Cong *et al*., 2021). Cong et al. (2021) have also suggested that PARPi resistance in BRCA1/53BP1 double-deficient cells is caused by restoration of the OFP backup pathway, evidenced by higher levels of chromatin-bound XRCC1 and LIG3 in these cells. While we find that the LIG3 BRCT domain, required for interaction with XRCC1, is critical for PARPi resistance in BRCA1/53BP1 double-deficient cells, we also find that PARPi-induced ssDNA gap formation in LIG3-depleted BRCA1/53BP1 double-deficient cells is fully rescued by MRE11 inhibition, indicating that LIG3 depletion in these cells does not impair OFP. Moreover, PARPi-induced ssDNA gaps in LIG3-depleted BRCA1/53BP1 double-deficient cells occur in both the newly replicated strands. Together, these data indicate that LIG3 is also involved in a separate, OFP- independent pathway of gap suppression.

Mechanistically, the LIG3-dependent gap suppression pathway might require repriming activities mediated by Polα, PRIMPOL or another unknown primase, for bypass of lesions such as PARPi-trapped PARP1 in BRCA1-deficient cells (García-Gómez *et al*., 2013; Fumasoni *et al*., 2015; Piberger *et al*., 2020; Quinet *et al*., 2020). These repriming activities could result in small gaps which require LIG3 to be filled. Loss of LIG3 in BRCA1-deficient and BRCA1/53BP1 double-deficient cells could thus result in the exposure of small ssDNA regions which would be a substrate for unscheduled MRE11-mediated processing. Subsequent processing of the small ssDNA regions could result in accumulation of longer stretches of post-replicative ssDNA, ultimately resulting in fork stalling, genomic instability and cell death (Figure 7).

PARP1 has recently been implicated in restraining RF speed in cells (Maya-Mendoza *et al*., 2018). We indeed observe an increase of fork speed in BRCA1-deficient cells treated with low doses of PARPi. Importantly, the increase in speed was specific to cells deficient for BRCA1, contrasting with the previous reports where PARP inhibition increased forks speed in BRCA1- proficient cells (Maya-Mendoza *et al*., 2018), possibly reflecting the use of higher olaparib concentrations and longer periods of exposure to PARPi in the latter study. In addition, we observe that PARPi treatment induces faster forks in PARPi-sensitive BRCA1-deficient cells as well as in PARPi-resistant BRCA/53BP1 double-deficient cells. Moreover, loss of LIG3 induces PARPi (hyper)sensitivity but suppresses PARPi-induced increase in fork speed in both BRCA1-deficient and BRCA1/53BP1 double-deficient cells. Together, these data show that PARPi-induced increase in fork speed in BRCA1-deficient cells is HR-independent and not causally related to PARPi sensitivity, in line with previous findings from Cong and colleagues (Cong *et al*., 2021) .

Our findings might have therapeutic implications, as LIG3 depletion also increases the efficacy of PARPi *in vivo*, resulting in prolonged survival of mice bearing PARPi-sensitive BRCA1- deficient or PARPi-resistant BRCA1/53BP1 double-deficient mammary tumors. Furthermore, we find LIG3 to be overexpressed in a portion of TNBC and serous ovarian cancers, further suggesting that LIG3 could possibly be targeted in these cancers. Pharmacological inhibition of LIG3 might therefore be a potential strategy to combat resistance to PARPi. Taken together, our findings establish loss of LIG3 as a potent enhancer of PARPi synthetic lethality in BRCA1-deficient tumors, irrespective of their HR status, and provide insights into the role of LIG3 in restraining replication stress and genome instability induced by BRCA1 loss.

## Limitations of this study

In this study we show that resistance to PARPi in BRCA1/53BP1 double-deficient cells is mediated by nuclear LIG3. As previously mentioned, mitochondrial LIG3 is essential for cellular viability and complete deletion of *Lig3* results in cellular death and early embryonic lethality in mice, whereas nuclear LIG3 is dispensable for cell viability (Simsek *et al*., 2011). In this study we have engineered BRCA1/53BP1 double-deficient mouse mammary tumor cells that only express mitochondrial *Lig3*, ensuring complete loss of nuclear *Lig3* expression. However, experiments testing olaparib sensitivity in other cell models were carried out using RNAi-mediated depletion which can result in downregulation of both isoforms. Thus, we cannot exclude the possibility that the observed effects are partially due to depletion of mitochondrial LIG3. In addition, our data indicate that loss of LIG3 has a more profound effect on PARPi sensitivity of BRCA1-deficient cells compared to BRCA2- deficient cells. However, it was not possible to compare the effects of LIG3 loss on PARPi sensitivity in isogenic cell lines deficient for either BRCA1 or BRCA2. Therefore, we cannot rule out that the observed differences were in part due to intercellular variability.

Although our findings might have clinical implications, datasets for large numbers of patients with *BRCA1*-mutated tumors who received PARPi treatment are not (yet) available. Finally, although our data suggest LIG3 as potential therapeutic target, small molecule inhibitors of LIG3 could target both nuclear and mitochondrial isoforms and might therefore result in undesirable toxicity.

## ACKNOWLEDGMENTS

We would like to thank Peter Bouwman and for kindly sharing the *Brca1^SCo/–^;Trp53^−/−^* mouse embryonic stem cells, Hanneke van der Gulden for technical assistance, Sylvie Noordermeer and Dan Durocher for kindly sharing the RPE1-TERT cells lines, and to Madalena Tarsounas for kindly sharing the DLD1 isogenic cell lines. We thank the members of the Preclinical Intervention Unit of the Mouse Clinic for Cancer and Ageing (MCCA) at the NKI for their technical support with the animal studies, and we thank the NKI core facilities: Digital Microscopy facility, Genomics Core facility and Animal facility for their excellent service. This work was supported by grants from the European Union’s Horizon 2020 research and innovation program under the Marie Skłodowska- Curie grant agreement No. 722729; the Dutch Research Council (NWO, VICI 91814643); the Danish Cancer Society (R204-A12617-B153); the Danish Council for Independent Research (DFF- 7016-00313); the Novo Nordisk Foundation (synergy grant no. 16854); the Swedish Research Council (VR-MH 2014-46602-117891-30); the Swedish Cancer Foundation (170176); the Danish National Research Foundation (project CARD, DNRF 125); the Dutch Cancer Society (KWF grant 11008 to ARC); and the Oncode Institute, which is partly financed by the KWF.

## AUTHOR CONTRIBUTIONS

Conceptualization – M.P.D., A.R.C. and J.J.; Methodology – M.P.D., I.v.d.H. and A.R.C.; Investigation – M.P.D., V.T., I.v.d.H., E.M., K.C., P.G., S.A., J. Bartkova, G.C.M.S. and M.A.S.; Supervision of *in vivo* experiments – M.v.d.V.; Data analysis – C.L., R.L.B, J.Bh. and S.Ch.; Writing of original draft, review & editing – M.P.D., A.R.C. and J.J.; Supervision – E.G., S.R., S.C., J. Bartek, A.R.C and J.J..

## STAR METHODS

### KEY RESOURCES TABLE

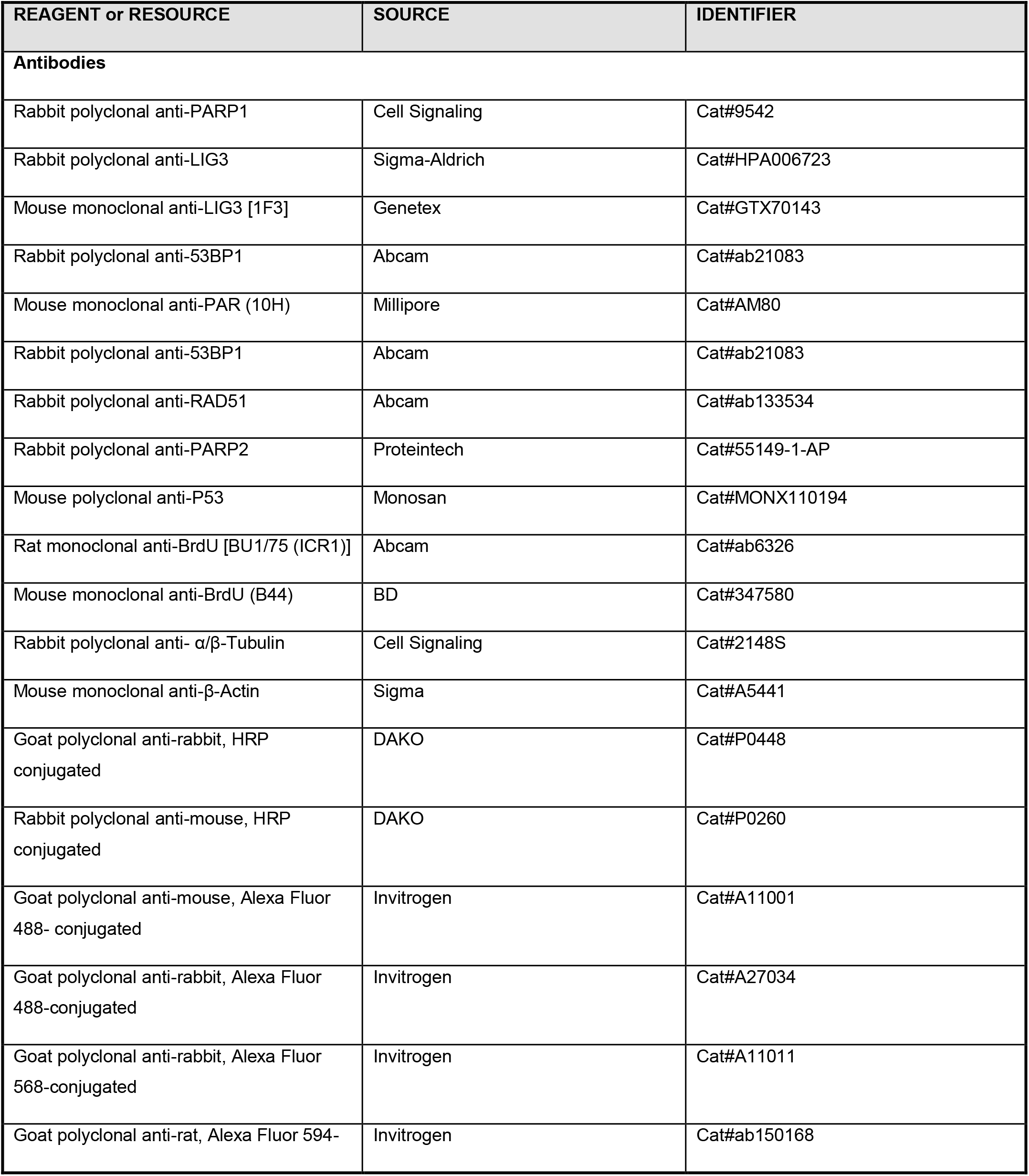
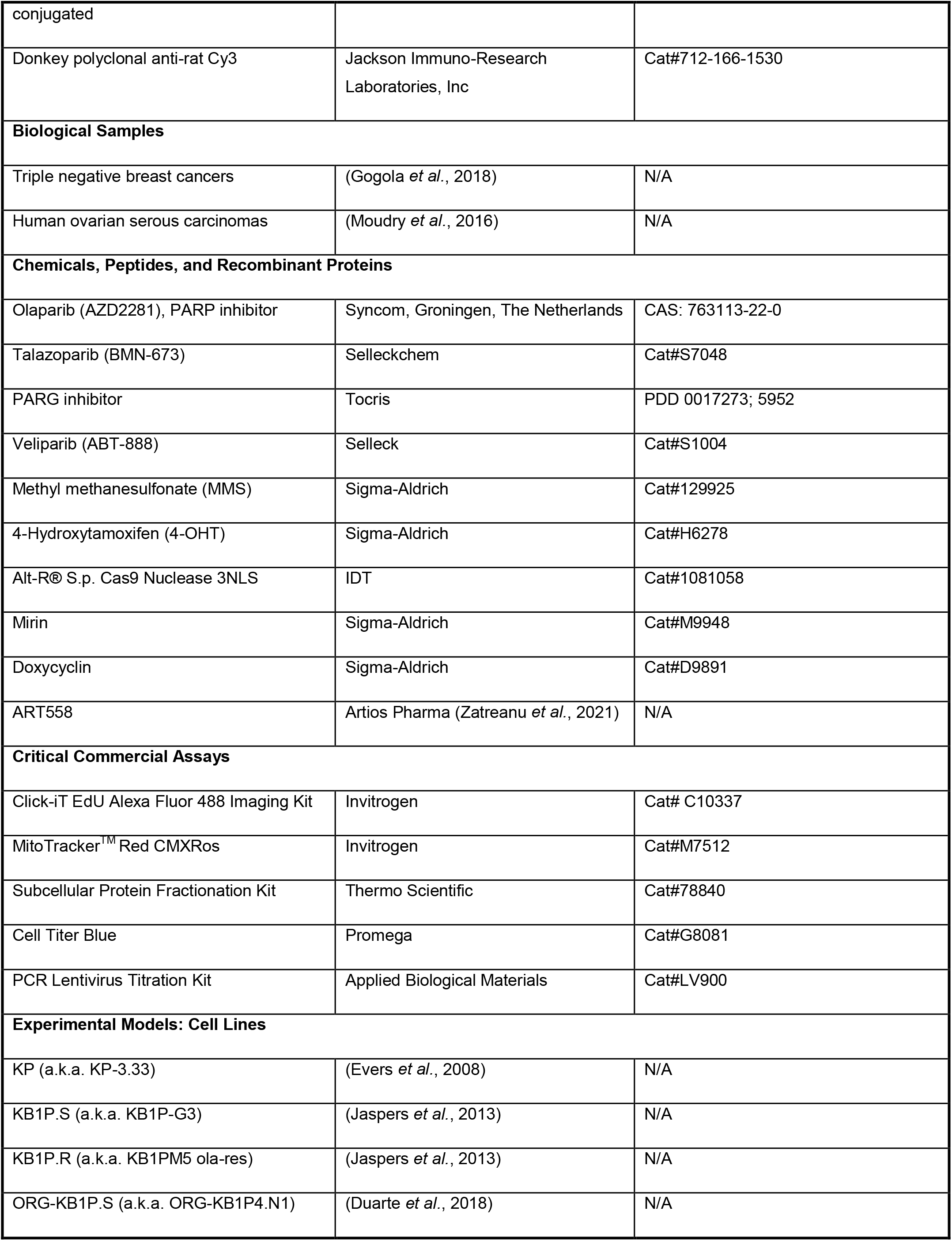
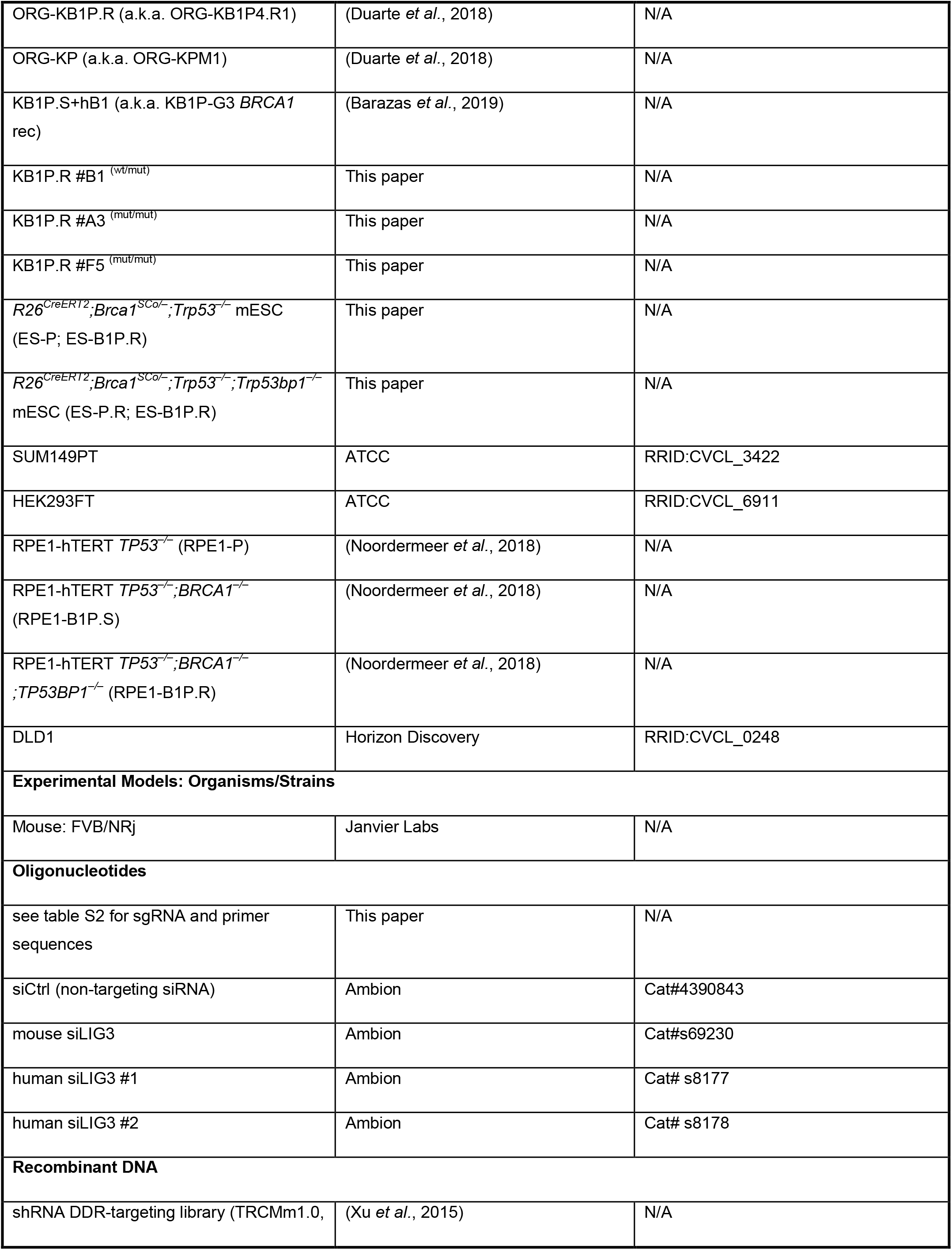
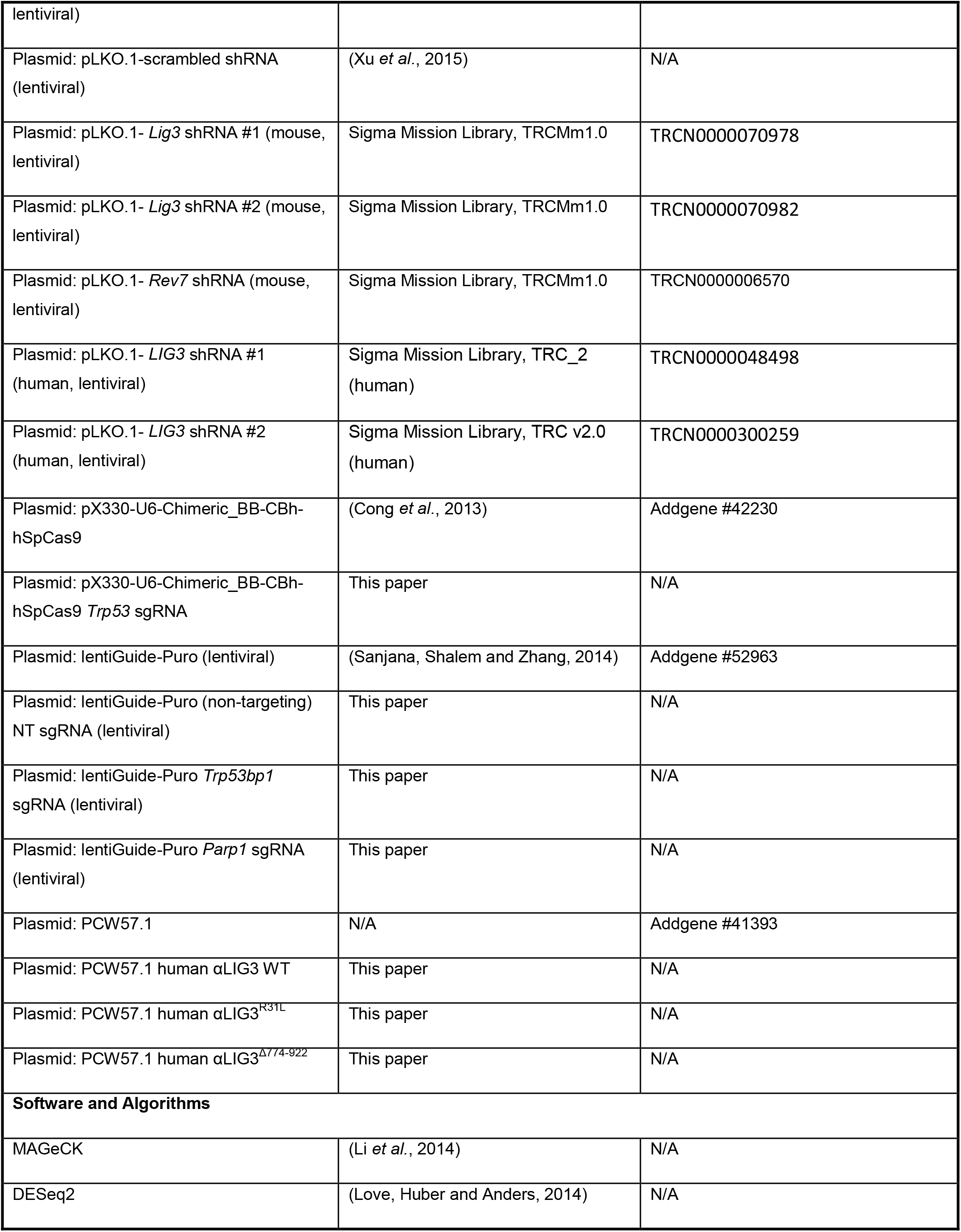
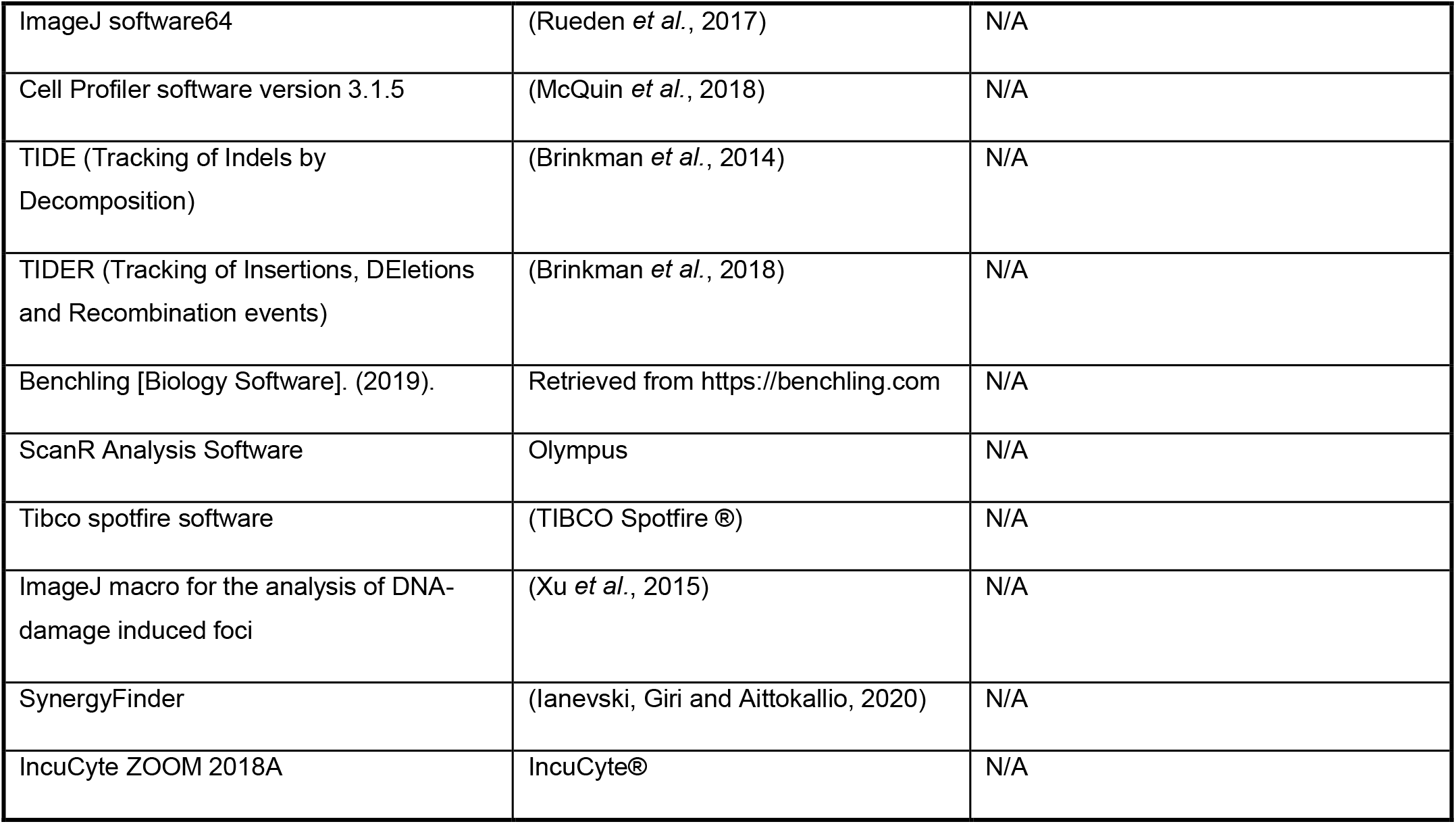

### CONTACT FOR REAGENT AND RESOURCE SHARING

Further information and requests for resources and reagents should be directed to and will be fulfilled by the Lead Contact, Jos Jonkers.

### EXPERIMENTAL MODEL AND SUBJECT DETAILS

#### Cell Lines

KP (Evers *et al*., 2008), KB1P.S, KB1P.R (Jaspers *et al*., 2013) and KB1P.S+hB1 (Barazas *et al*., 2019) have been previously described. LIG3 nuclear mutants, KB1P.R-B1, KB1P.R-A3 and KB1P.R-F5, have been generated in this study. All these cell lines were cultured in in DMEM/F12+GlutaMAX (Gibco) containing 5μg/ml Insulin (Sigma, #I0516), 5 ng/ml cholera toxin (Sigma, #C8052), 5 ng/ml murine epidermal growth-factor (EGF, Sigma, #E4127), 10% FBS and 50 units/ml penicillin-streptomycin (Gibco) and were cultured under low oxygen conditions (3% O2, 5% CO_2_ at 37°C). Mouse ES cells with a selectable conditional *Brca1* deletion (*R26Cre^ERT2/wt^;Brca1^SCo/–^*) have been previously described (Bouwman et al. 2010). Additional knockout of *Trp53, Trp53bp1* and *Parp1* has been generated in this study. These cells were cultured on gelatin-coated plates in 60% buffalo red liver (BRL) cell conditioned medium, 0.1 mM β-mercaptoethanol (Merck) and 10^3^

U/ml ESGRO LIF (Millipore) and 50 units/ml penicillin-streptomycin (Gibco) under normal oxygen conditions (21% O2, 5% CO_2_, 37°C). SUM149PT (RRID: CVCL_3422) cells were grown in RPMI1640 (Gibco) medium supplied with 10% fetal calf serum and 50 units/ml penicillin-streptomycin (Gibco). RPE1-hTERT and DLD-1 cell lines were grown in DMEM+GlutaMAX (Gibco) supplemented with 10% FBS and 50 units/ml penicillin-streptomycin (Gibco). RPE1-P, RPE1-B1P.S and RPE1-B1P.R cells were generated by Noordermeer et al. 2018. HEK293FT (RRID: CVCL_6911) cells were cultured in IMDM+GlutaMAX-I (Gibco) supplemented with 10% FBS and 50 units/ml penicillin-streptomycin (Gibco). SUM149PT and DLD1 cell lines were cultured under normal oxygen conditions (21% O_2_, 5% CO_2_, 37°C). RPE1 cell lines were cultured under low oxygen conditions (3% O_2_, 5% CO_2_ at 37°C).

#### Tumor-Derived Organoids

All lines have been described before (Duarte *et al*., 2018). ORG-KB1P.S and ORG-KB1P.R tumor organoids were derived from a PARPi-naïve and PARPi-resistant *K14cre;Brca1^F/F/^;Trp53^F/F^* (KB1P) mouse mammary tumor, respectively. The ORG-KP tumor organoid line was derived from a *K14cre;Trp53^F/F^;Abcb1a^−/−^;Abcb1b^−/−^* (KPM) mouse mammary tumor. Cultures were embedded in Culturex Reduced Growth Factor Basement Membrane Extract Type 2 (BME, Trevigen; 40 ml BME:growth media 1:1 drop in a single well of 24-well plate) and grown in Advanced DMEM/F12 (Gibco) supplemented with 1M HEPES (Gibco), GlutaMAX (Gibco), 50 units/ml penicillin-streptomycin (Gibco), B27 (Gibco), 125 mM N-acetyl-L-cysteine (Sigma) and 50 ng/ml murine epidermal growth factor (Sigma). Organoids were cultured under standard conditions (37°C, 5% CO_2_) and regularly tested for mycoplasma contamination.

#### Mice

All animal experiments were approved by the Animal Ethics Committee of The Netherlands Cancer Institute (Amsterdam, the Netherlands) and performed in accordance with the Dutch Act on Animal Experimentation (November 2014). Organoid transplantation experiments were performed in syngeneic, wild-type F1 FVB (FVB/NRj) females, at the age of 6 weeks. Parental FVB animals were purchased from Janvier Labs. Animals were assigned randomly to the treatment groups and the treatments were supported by animal technicians who were blinded regarding the hypothesis of the treatment outcome.

#### Human Samples of TNBC and Ovarian Serous Carcinomas

Samples were previously described in (Gogola *et al*., 2018). Retrospective Triple Negative Breast Cancer (TNBCs) biopsies from 86 clinical high-risk patients (high-risk definition according to the Danish Breast Cooperative Group (www.dbcg.dk accessed 22.10.2009) that underwent mastectomy between 2003 and 2015 were selected and classified as being triple negative according to the criteria set in the ASCO/CAP guidelines (ER<1%, PR<1%, HER2 0, 1+ or 2+ but FISH/ CISH negative). The patients presented a unifocal tumor of an estimated size of more than 20 mm. None of the patients had previous surgery to the breast and did not receive preoperative treatment. This study was conducted in compliance with the Helsinki II Declaration and written informed consent was obtained from all participants and approved by the Copenhagen and Frederiksberg regional division of the Danish National Committee on Biomedical Research Ethics (KF 01-069/03). Paraffin- embedded material from the cohort of ovarian tumors was collected at the Department of Pathology, University Hospital, Las Palmas, Gran Canaria, Spain, from surgical operations performed in the period 1995-2005. For the purpose of the present study, only samples from serous ovarian carcinoma (the type approved for treatment by PARP inhibitors) were used from a larger cohort that was reported previously (Moudry *et al*., 2016), and included also other histological types of ovarian tumors. The use of long-term stored tissue samples in this study was in accordance with the Spanish codes of conduct (Ley de Investigación Biomédica) and was approved by the review board of the participating institution. Patients were informed that samples may be used for research purposes under the premise of anonymity.

### METHOD DETAILS

#### Functional Genetic Screens

The DDR shRNA library was stably introduced into *Brca1^−/−^;Trp53^−/−^;Trp53bp1^−/−^* mESCs and in KB1P4.R1 by lentiviral transduction using a multiplicity of transduction (MOI) of 1, in order to ensure that each cell only gets incorporated with one only sgRNA. mES cells and organoids were selected with puromycin, 3 μg/ml, for 3 days and then seeded in the presence of PARPi (IC50<30, mES cells, 25nM olaparib; organoids, 50nM), left untreated or pelleted for the genomic DNA isolation (T0). The total number of cells used in a single screen was calculated as following: library complexity x coverage (5000x in mESc, 1000x in organoids). Cells were kept in culture for 3 weeks and passaged every 5 days (and seeded in single cells) while keeping the coverage at every passage. mES cells were seeded at a density of 2,500 cells per 15 cm dish and organoids at a density of 50,000 cells/well, 24-well format. Screens were done in triplicate for each condition. In the end of the screen, cells were pooled and genomic DNA was extracted (QIAmp DNA Mini Kit, Qiagen). shRNA sequences were retrieved by a two-step PCR amplification, as described before (Xu *et al*., 2015). To maintain screening coverage, the amount of genomic DNA used as an input for the first PCR reaction was taken into account (6 μg of genomic DNA per 10^6^ genomes, 1 μg/PCR reaction). Resulting PCR products were purified using MiniElute PCR Purification Kit (Qiagen) and submitted for Illumina sequencing. Sequence alignment and dropout analysis was carried out using the algorithms MAGeCK (Li et al., 2014) (FDR <= 0.1) and DESeq2 (Love, Huber and Anders, 2014) (FDR <= 0.05, log2Fc <=-2, baseMean >= 100, at least 3 hit shRNA in the depletion direction and none in the opposite direction). In order to reduce the noise level, we filtered out sgRNAs with low counts in the T0 sample: mESc, sum of the three T0 samples >= 10, organoids, mean over the three T0 samples >= 50. Gene ranking is generated automatically with MaGECK algorithm. To generate gene ranking based on DESeq2 algorithm, we calculated per gene the number of hit shRNAs and the mean of the log2FoldChange over those shRNAs. We then ranked the genes based on these two metrics.

#### Constructs

A collection of 1,976 lentiviral hairpins targeting 391 DDR-related mouse genes (pLKO.1; DDR library) was derived from the Sigma Mission library (TRCMm1.0) as described before (Xu *et al*., 2015). Individual hairpin constructs used in the validation studies were selected from the TRC library: mouse LIG3 shRNA #1: TRCN0000070978, mouse LIG3 shRNA #2: TRCN0000070982, mouse REV7 shRNA: TRCN0000006570, human LIG3 shRNA #1: TRCN0000048498, human LIG3 shRNA #2: TRCN0000300259. For CRISPR/Cas9-mediated genome editing of *Parp1*, a sgRNAs was cloned into plentiGuide-Puro (lentiviral) as described previously (Sanjana, Shalem and Zhang, 2014). For the LIG3 overexpression constructs, human α-LIG3 wild type, human α-LIG3 carrying a mutation in the PARP-like ZnF domain (R31L), and human α-LIG3 with a C-terminal Δ774-922 truncation which includes the BRCT domain were cloned into PCW57.1 plasmid. All constructs were verified by Sanger sequencing.

#### Lentiviral Transductions

Lentiviral stocks, pseudotyped with the VSV-G envelope, were generated by transient transfection of HEK293FT cells, as described before (Follenzi *et al*., 2000). Production of integration-deficient lentivirus (IDLV) stocks was carried out in a similar fashion, with the exception that the packaging plasmid contains a point mutation in the integrase gene (psPAX2, gift from Bastian Evers). Lentiviral titers were determined using the qPCR Lentivirus Titration Kit (Applied Biological Materials), following the manufacturer’s instructions. For all experiments the amount of lentiviral supernatant used was calculated to achieve an MOI of 50, except for the transduction of the lentiviral library for which a MOI of 1 was used, as described above. Cells were incubated with lentiviral supernatants overnight in the presence of polybrene (8 μg/ml). Tumor-derived organoids were transduced according to a previously established protocol (Duarte *et al*., 2018). Antibiotic selection was initiated right after transduction for cells, 24h after transduction in organoids, and was carried out for 3 consecutive days.

#### Genome Editing

For CRISPR/Cas9-mediated genome editing of *Trp53* in mESCs, *R26Cre^ERT2/wt^;Brca1^SCo/–^* cells (Bouwman *et al*., 2010) were transiently transfected with a modified a pX330-U6-Chimeric-BB-CBh- hSpCas9 plasmid containing a puromycin resistance marker (Cong *et al*., 2013; Drost *et al*., 2016) in which a sgRNA targeting *Trp53* was cloned. Knockout clones were selected under puromycin for 3 days and tested by TIDE and western blot.

For CRISPR/Cas9-mediated genome editing of *Trp53bp1* in mESCs, Cas9-expressing *R26Cre^ERT2/Cas9^;Brca1^SCo/–^;Trp53^−/−^ cells* (Barazas *et al*., 2018) were incubated with lentiviral supernatants of pLentiGuide-Puro cloned with a sgRNA targeting *Trp53bp1.* After selection with puromycin for 3 days, surviving cells were subcloned and tested by TIDE and western blot.

For CRISPR/Cas9-mediated genome editing of *Parp1, the* Cas9-expressing *R26Cre^ERT2/Cas9^; Brca1^−/−^;Trp53^−/−^;Trp53bp1^−/−^* mESCs were incubated with lentiviral supernatants of pLentiGuide-Puro cloned with a sgRNA targeting *Parp1*. After selection with puromycin for 3 days, surviving cells were subcloned and tested by TIDE and western blot.

For the disruption of the starting codon encoding for nuclear LIG3, the desired mutation (ATG>CTC) was introduced in KB1P.R mouse tumor cells according to the Alt-R CRISPR-Cas9 System of IDT (Yoshimi *et al*., 2016). Briefly, the crRNA targeting sequence and the homology template, a 120bp ssODN, were designed using CRISPR design tools of Benchling. While the sgRNA was designed to target the nuclear ATG, the homology template contains an ATG>CTC mutation, encoding a leucine instead of the original methionine. 10 µl tracrRNA (100 µM) and 10 µl crRNA (100 µM) were annealed in 80 μl nuclease free duplex buffer (IDT#11-05-01-03) to form a 10µM gRNA solution. The ssODN template was also annealed to form a 10µM solution. 6 μl of 10 µM sgRNA, 6 µl of 10 µM Cas9 protein, and 6 µl of 10 µM ssODN (Ultramer IDT) were mixed in optiMEM (Gibco), to final volume of 125 µl and incubated for 5 min at RT (Mix 1). Then, 3µl of Lipofectamine RNAiMAX (Invitrogen) were mixed with 122 µl with optiMEM (Mix 2). Mix 1 and mix 2 were mixed together and incubated at RT for 20 min. During these 20 min, 150.000 cells were trypsinized and collected in 750 µl of medium. The 250 µl Mix was then added to the cells in a 12-well for reverse transfection. Next day cells were expanded and 3 days after transfection the cells were harvested for analysis of the genomic DNA.

To assess modification rate, genomic DNA was extracted (Puregene Core Kit A, Qiagen) and 100 ng was used as an input for the PCR amplification of the targeted sequence. PCR reaction was performed with Thermo Scientific Phusion High-Fidelity PCR Master Mix (Thermo Scientific), according to manufacturer’s instructions (3-step protocol: annealing - 60C for 5 s, extension time 30 s) and using primers listed in Table S2. Resulting PCR products served as a template for the BigDye Terminator v3.1 reaction (Thermo Fisher). BigDye PCR reactions were performed with the same forward primers as in the preceding PCR reactions (no reverse primer used) and according to the BigDye manufacturer’s protocol. For knockout, allele composition was determined with the TIDE analysis (Brinkman *et al*., 2014) by comparing sequences from modified and parental (transduced with non-targeting sgRNAs) cells. For knock-in, allele composition was determined with the TIDER analysis (Brinkman *et al*., 2018) by comparing sequences from modified and parental cells (transduced with non-targeting sgRNAs), and reference template. The later was generated with a simple two-step PCR protocol, with two complementary primers designed to carry the designed mutations as present in the donor template (Brinkman et al. 2018).

#### siRNA and Transfections

Non-targeting siRNA and siRNA against mouse and human LIG3 were transfected into the cells using Lipofectamine RNAiMAX (Invitrogen) according to the manufacturer’s instructions. All experiments were carried out between 48 and 72hr post-transfection.

#### Long-Term Clonogenic Assays

Long-term clonogenic assays were always performed in 6-well plates, with exception of organoids which were cultured in 24-well plated as described before, and to DLD-1 cells which was performed in a 12-well plate. Cells were seeded at low density to avoid contact inhibition between the clones (KB1P.S: 5,000 cells/well; KB1P.R: 2,500 cells/well; ORG-KB1P.S and ORG-KB1P.R: 50.000 cells/well; ES-B1P.R and ES-P.R: 3,000 cells/well; ES-B1P.S and ES-P: 5,000 cells/well; SUM149PT: 5,000 cells/well; RPE1-P: 3,000 cells/well, RPE1-B1P.S and RPE1-B1P.R: 5,000 cells/well; DLD-1: 3,500 cells/well; DLD-1 BRCA2 KO cells: 5,000) and cultured for 10-15 days. Media was refreshed once a week. For the quantification, cells were incubated with Cell-Titer Blue (Promega) reagent and later fixed with 4% formaldehyde and stained with 0.1% crystal violet. Drug treatments: cells were grown in the continuous presence of PARPi (olaparib, talazoparib or veliparib) at the indicated concentrations. mESCs with a selectable conditional *Brca1* deletion were treated with 0.5µM 4OHT for 3 days right before the start of the clonogenic assay, when indicated. PARPis were reconstituted in DMSO (10 mM) and 4OHT in EtOH (2.5 mM). Expression of human LIG3 constructs was induced with treatment with 2μM Doxycycline for two days prior to the start of the assay and at the start of the assay.

#### Proliferation assay

Cell were imaged every 4h using IncuCyte ®, for 1 week duration. Cells were seeded at low density and grown under normal oxygen conditions (21% O_2_, 5% CO_2_, 37°C). Data was analyzed using IncuCyte ZOOM 2018A software.

#### RT-qPCR

In order to determine gene expression levels, RNA was extracted from cultured cells using ISOLATE II RNA Mini Kit (Bioline) and used as a template to generate cDNA with Tetro cDNA Synthesis Kit (Bioline). Quantitative RT-PCR was performed using SensiMix SYBR Low-ROX Kit (Bioline; annealing temperature – 60°C) in a Lightcycler 480 384-well plate (Roche), and analyzed using Lightcycler 480 Software v1.5 (Roche). Mouse *Rps20* and human *HPRT* were used as house- keeping genes. The primer sequences used in this study are listed in Table S2.

#### Western Blot

Cells were trypsinized and then lysed in lysis buffer (20 mM Tris pH 8.0, 300 mM NaCl, 2% NP40, 20% glycerol, 10 mM EDTA, protease inhibitors (cOmplete Mini EDTA-free, Roche, 100x stock)), for 20 min. For PAR detection in PARP1 knockout mES cells, 10µM PARGi was added to the lysis buffer, when indicated. For P53 detection, cells were irradiated at 15 x 100 μJ/cm^2^. The protein concentration was determined using Pierce BCA Protein Assay Kit (Thermo Scientific). SDS-Page was carried out with the Invitrogen NuPAGE SDS-PAGE Gel System (Thermo Fisher; for LIG3: 2- 8% Tris-acetate gels were used, buffer Tris-Acetate; for all other proteins: 4–12% Bis-Tris gels were used, buffer: MOPS; input: 40µg protein), according to the manufacturer’s protocol. Next, proteins were electrophoretically transferred to a nitrocellulose membrane (Biorad). Before blocking, membranes were stained with Ponceau S, followed by blocking in 5% (w/v) milk in TBS-T for 1hr at RT. Membranes were incubated with primary antibody 4hrs at RT in 1% (w/v) milk in TBS-T (rabbit anti-PARP1, 1:1000; rabbit anti-H3, 1:5000; mouse anti-lig3, 1:500; rabbit anti-lig3, 1:1000; rabbit anti-tubulin, 1:1000; anti-PAR, 1:1000; mouse anti-P53, 1:1000). Horseradish peroxidase (HRP)- con-jugated secondary antibody incubation was performed for 1 hr at RT (anti-mouse or anti-rabbit HRP 1:2000) in 1% (w/v) milk in TBS-T. Signals were visualized by ECL (Pierce ECL Western Blotting Substrate, Thermo Scientific).

#### Cytotoxicity Assays

Cytotoxicity assays were carried in 96-well plates, for 3 days. Olaparib and POLθ inhibitor ART558 were used at the indicated concentrations. Olaparib was used at concentrations that wouldn’t lead to lethality of LIG3-depleted cells when used as single agent in order to allow a window to detect the effect of POLθ inhibition. KB1P.R and KB1P.R A3 cells were seeded at high density, 2.000 cells/well. For the quantification, cells were incubated with Cell-Titer Blue (Promega) reagent. The expected drug combination responses were calculated based on Bliss reference model using SynergyFinder (Ianevski, Giri and Aittokallio, 2020).

#### Proximity ligation assay (PLA)

Protocol was carried out as mentioned previously (Mukherjee *et al*., 2019). On coverslips, cells were grown to a confluence of 60-70%. On the day of the experiment, cells were incubated with PARGi (10µM) for a total of 30 minutes or 0.5μM olaparib for 2hr and the final 10 minutes cells were incubated with EdU (20μM) during PARGi incubation to visualize S-phase cells. After EdU labeling cells were gently washed two times with PBS and fixed with 4% paraformaldehyde for 15 min at RT. PFA was discarded after fixation and slides were washed with cold PBS for 8 minutes each three times. Cells were next permeabilized by incubating the coverslips in PBS containing 0.5% Triton-X for 15 min at RT and subsequently washed in PBS twice for 5 min each. Freshly prepared click reaction mix (2mM of copper sulfate, 10 μM of biotin-azide and 100 mM of sodium ascorbate were added to PBS in that order and mixed well) was applied to the slides (30 μl/slide) in a humid chamber and incubated for 1 hr at RT. Slides were washed with PBS for 5 min after the click reaction and placed back in the humid chamber and blocked at room temperature for 1 hr with a blocking buffer (10% goat serum and 0.1%Triton X-100 in PBS). In combination with anti-biotin (1:1000), rabbit anti-LIG3 (1:150, Sigma-Aldrich, #HPA006723) primary antibody was diluted in a blocking solution, dispensed to slides (30 μl/slide) and incubated in a humid chamber at 4°C overnight. Slides were washed with wash buffer A (0.01 M Tris-HCl, 0.15 M NaCl, and 0.05 % Tween-20, pH 7.4) for 5 min each after overnight incubation. Duolink In Situ PLA probes, the anti- mouse plus and anti-rabbit minus were diluted 1:5 in the blocking solution (10% goat serum and 0.1% Triton X-100 in PBS), dispensed to slides (30 μl/well) and incubated at 37°C for 1 hr. Slides were washed three times with buffer-A, 5 min each. The ligation mix was prepared by diluting Duolink ligation stock (1:5) and ligase (1:40) in high purity water and was applied to slides (30 μl/well) and incubated at 37°C for 30 min. Slides were washed with buffer-A twice for 2 min each. Amplification mix was prepared by diluting Duolink amplification stock (1:5) and rolling circle polymerase (1:80) in high-purity water and applied to slides (30μl /well) and incubated for 100 min at 37°C in a humid chamber. Slides were washed with wash buffer-B solution (0.2 M Tris and 0.1 M NaCl) three times for 10 min each and one time in 0.01X diluted wash buffer-B solution for 1 min. Coverslips were incubated with DAPI for 5 min and mounted with ProLong Gold antifade reagent (Invitrogen) and imaged using confocal and analyzed using ImageJ software 64.

#### Immunofluorescence

##### RAD51 IRIF

Cells were seeded on Millicell EZ slides (#PEZGS0816, Millipore) 24 hr prior the assay to achieve ∼90% confluency. Cells were then irradiated using the Gammacell 40 Extractor (Best Theratronics Ltd.) at the dose of 10 Gy and allowed to recover for 3 hr. Cells washed with PBS++ (PBS solution containing 1 mM CaCl2 and 0.5 mM MgCl2) and pre-extracted with 0.5% (v/v) Triton X-100 in PBS++ for 5 min. Next, cells were washed with PBS++ and fixed with 2% (v/v) paraformaldehyde solution in PBS for 20 min. Next, cells were permeabilized with ice-cold methanol/acetone solution (1:1) for 15 min. To minimize the background, cells were further incubated for 20 min in staining buffer (1% (w/v) BSA, 1% (v/v) FBS, 0.15% (w/v) glycine and 0.1% (v/v) Triton X-100 in PBS). Staining buffer was also used as a solvent for antibodies – primary antibody anti-RAD51, 1:1500, #ab133534, abcam; secondary antibody Alexa Fluor® 658-conjugated, 1:1000, A11011, Invitrogen. Incubation with primary and secondary antibodies was done for 2 hr and 1 hr, respectively. All incubations were performed at room temperature. Samples were mounted with VECTASHIELD Hard Set Mounting Media with DAPI (#H-1500; Vector Laboratories). Images were captured with Leica SP5 (Leica Microsystems) confocal system and analyzed using an in-house developed macro to automatically and objectively evaluate the DNA damage-induced foci (Xu *et al*., 2015). As a positive and negative control for RAD51 staining, BRCA-proficient KP and BRCA1-deficient KB1P.S cells were used.

##### LIG3-EdU co-localization

Cells were incubated with 20 µM EdU for 1hr to visualize cells in S-phase. In the last 20 min, 10µM PARGi was added to the medium. Cells washed with PBS and pre-extracted with CSK50 buffer for 7 min (10µM PARGi PDDX-001 was added to pre-extraction buffer). Cells were washed with PBS and fixed with 4% formaldehyde, followed by three washes with PBS and permeabilization with ice- cold methanol/acetone solution (1:1). EdU Click-iT reaction mix was added to each well and incubated at RT for 30 min. Fixed cells were washed three times with staining buffer (5% (v/v) FBS, 5% (w/v) BSA, and 0.05% (v/v) Tween-20 in PBS) and incubated with primary antibody anti-LIG3 (1:150, Sigma-Aldrich, #HPA006723) in staining buffer for 2hr at RT. After three washes in staining buffer, cells were incubated with secondary antibody anti–rabbit Alexa Fluor 488 (1:500, A27034, Invitrogen) in staining buffer, followed by three last washes in staining buffer and one wash in PBS. Samples were mounted with VECTASHIELD Hard Set Mounting Media with DAPI (#H-1500; Vector Laboratories). Images were captured with Leica SP5 (Leica Microsystems) confocal system and analyzed with ImageJ software.

##### Native BrdU

Cells were labeled with 10μM BrdU for 48hr. When indicated, cells were incubated with Mirin (25μM) for the same 48hr. Upon treatment with the final 2hr PARPi inhibitor (0.5μM), the cells were washed with PBS and pre-extracted by CSK-buffer (PIPES 10mM, NaCl 100mM, Sucrose 300mM, EGTA 250mM, MgCl2 1mM, DTT 1mM and protease inhibitors cocktail) on ice for 5 minutes. Cells were then fixed using 4% formaldehyde (FA) for 15 min at RT, and then permeabilized by 0.5% Triton X-100 in CSK-buffer. Permeabilized cells were then incubated with primary antibody against anti-BrdU antibody (Abcam 6326) at 37°C for 1 hr. Cells were washed and incubated with secondary antibodies (Alexa Fluor 594) for 1h at room temp. After the wash cells were incubated with DAPI (0.1μg/ml) for 5 minutes. For mouse tumor cells (high content imaging), DAPI and ssDNA signal, Z-stack of 6 stacks (1mm/stack) covering at least 75 fields were imaged. Results were analyzed using DAPI channel and filtered with roundness and size of the nucleus. The quantification of pixel intensities (mean, median and sum) for each nucleus was calculated in the DAPI and 594 nm channels. The quantified values obtained were exported to Tibco spotfire software (TIBCO Spotfire ®) for the generation of scatter plots. For human RPE1 cells, images were collected by fluorescence microscopy (Axioplan 2 and Axio Observer, Zeiss) at a constant exposure time in each experiment. Representative images were processed by ImageJ software. Mean intensities of ssDNA in each nucleus were measured with Cell Profiler software version 3.1.5 from Broad Institute.

#### DNA Fiber assay

##### Mouse tumor cells

DNA fiber analysis was conducted in accordance with the previously described protocol (Ray Chaudhuri *et al*., 2012). Briefly, cells were transfected for 48 hours followed by treatment with olaparib (0.5µM), or left untreated, for the final two hours. Cells were sequentially pulse-labelled with nucleotide analogues, 30µM CldU (c6891, Sigma-Aldrich) and 250µM IdU (I0050000, European Pharmacopoeia) for 20 min during the incubation of olaparib. After double labelling, cells were washed with PBS, harvested and resuspended in ice cold PBS to the final concentration 2.5 × 105 cells per ml. Labelled cells were mixed with unlabeled cells at 1:1 (v/v), and 2.5 µl of cell suspension was spotted at the end of the microscope slide. 8 µl of lysis buffer (200mM Tris-HCl, pH 7.5, 50mM EDTA, and 0.5% (w/v) SDS) was applied on the top of the cell suspension, then mixed by gently stirring with the pipette tip and incubated for 8 min. Following cell lysis, slides were tilted to 15–45° to allow the DNA fibers spreading along the slide, air dried, fixed in 3:1 methanol/acetic acid overnight at 4 °C. Subsequently, fibers were denatured with 2.5 M HCl for 1 hr. After denaturation, slides were washed with PBS and blocked in blocking solution (0.2% Tween 20 in 1% BSA/PBS) for 40 min. After blocking, primary antibody solutions are applied, anti-BrdU antibody recognizing CldU (1:500, ab6326; Abcam) and IdU (1:100, B44, 347580; BD) for 2 hours in the dark at RT followed by 1h incubation with secondary antibodies: anti–mouse Alexa Fluor 488 (1:300, A11001, Invitrogen) and anti–rat Cy3 (1:150, 712-166-153, Jackson Immuno-Research Laboratories, Inc.). Finally, slides are washed with PBS and subsequently mounting medium is spotted and coverslips are applied by gently pressing down. Slides were sealed with nail polish and air dried. Fibers were visualized and imaged by Carl Zeiis Axio Imager D2 microscope using 63X Plan Apo1.4 NA oil immersion objective. Data analysis was carried out with ImageJ software64.

##### RPE1-hTERT cells

These assays were performed as previously described (Peng *et al*., 2018; Cong *et al*., 2021). Briefly, cells were treated for 2 hr with 0.5µM olaparib or left untreated. During the last 40 min, cells were labeled by sequential incorporation of IdU and CldU into nascent DNA strand. Cells were then collected, washed, spotted, and lysed on positively charged microscope slides by 7.5 mL spreading buffer for 8 min at RT. Individual DNA fibers were released and spread by tilting the slides at 45°C. After air-drying, fibers were fixed by 3:1 methanol/acetic acid at RT for 3 min. Fibers were then rehydrated in PBS, denatured with 2.5 M HCl for 30 min, washed with PBS, and blocked with blocking buffer for 1 hr. Next, slides were incubated for 2.5 hr with primary antibodies diluted in blocking buffer (IdU, B44, 347580; BD; CldU, ab6326, Abcam), washed several times in PBS, and then incubated with secondary antibodies in blocking buffer for 1 hr (IdU, goat anti-mouse, Alexa 488; CldU, goat anti-rat, Alexa Fluor 594). After washing and air-drying, slides were mounted with Prolong (Invitrogen, P36930). Finally, fibers were visualized and imaged with Axioplan 2 imaging, Zeiss.

### Metaphase spreads and telomere FISH

Metaphase spreads were carried out according to the standard protocol described previously (Mukherjee *et al*., 2019). Briefly, exponentially growing cells (50–80 % confluence) were treated with 0.5µM olaparib for 2hr or left untreated, and recovered for 6 hr. Post treatment, drug treated medium was washed out and cells were allowed to grow in complete growth medium and exposed with colcemid for 8 h. Metaphase spreads were prepared by conventional methods and check under the microscope before telomere labelling. Metaphase slides in coplin jar containing 2X SSC buffer (Sigma-S6639) were equilibrated at room temperature for 10 minutes. Proteins were digested by incubation of the slides in pre-warmed 0.01M HCl containing pepsin for 1.5 min at 37°C. Slides were washed twice with PBS 5 min each and then one time with 1 M MgCl2 in 1X PBS for 5 min. After washing slides were placed in coplin jar containing 1% formaldehyde and fixed for 10 mins at RT without shaking. Slides were washed with PBS and dehydrated in the ethanol series: 70%, 90% and 100% for 3 minutes each and air dried. Next, slides were denatured in 70% deionized formamide at 80°C for 1 min 15 sec and immediately placed in chilled ethanol series 70%, 90% and 100% for 3 minutes each and allowed slides for air dry. Pre-annealed telomere probes were added to the denatured slides and allowed for hybridization at 37°C in hybridization chamber for 40 minutes. After hybridization slides were washed sequentially 3 times each with 50% formamide in 2X SSC (preheated to 45°C), 0.1X SSC (preheated to 60°C), 4X SSC (0.1% Tween-20), and 2X SSC respectively. Slides were allowed to air dry and mounted using DAPI anti-fade. A minimum 60 metaphase images were captured using Carl Zeiss Axio Imager D2 microscope using 63x Plan Apo 1.4 NA oil immersion objective and analyzed with ImageJ software64 for chromosomal aberrations.

#### Electron microscopy analysis

EM analysis was performed according to the standard protocol (Zellweger *et al*., 2015). For DNA extraction, cells were lysed in lysis buffer and digested at 50 °C in the presence of Proteinase-K for 2hr. The DNA was purified using chloroform/isoamyl alcohol and precipitated in isopropanol and given 70% ethanol wash and resuspended in elution buffer (TE). Isolated genomic DNA was digested with PvuII HF restriction enzyme for 4 to 5 hr. Replication intermediates were enriched by using QIAGEN G-100 columns (as manufacture’s protocol) and concentrated by an Amicon size- exclusion column. The benzyldimethylalkylammonium chloride (BAC) method was used to spread the DNA on the water surface and then loaded on carbon-coated nickel grids and finally DNA was coated with platinum using high-vacuum evaporator MED 010 (Bal Tec). Microscopy was performed with a transmission electron microscope FEI Talos, with 4 K by 4 K cmos camera. For each experimental condition, at least 70 RF intermediates were analyzed per experiment and ImageJ software64 was used to process analyze the images.

#### DSB detection by PFGE

DSB detection by PFGE was done as reported previously (Cornacchia *et al*., 2012). Cells were casted into 0.8% agarose plugs (2.5 x 105 cells/plug), digested in lysis buffer (100 mM EDTA, 1% sodium lauryl sarcosine, 0.2% sodium deoxycholate, 1 mg/ml proteinase-K) at 37 °C for 36–40 h, and washed in 10 mM Tris-HCl (pH 8.0)–100 mM EDTA. Electrophoresis was performed at 14 °C in 0.9% pulse field-certified agarose (Bio-Rad) using Tris-borate-EDTA buffer in a Bio-Rad Chef DR III apparatus (9 h, 120°, 5.5 V/cm, and 30- to 18-s switch time; 6 h, 117°, 4.5 V/cm, and 18- to 9-s switch time; and 6 h, 112°, 4 V/cm, and 9- to 5-s switch time). The gel was stained with ethidium bromide and imaged on Uvidoc-HD2 Imager. Quantification of DSB was carried out using ImageJ software64. Relative DSB levels were calculated by comparing the results in the treatment conditions to that of the DSB level observed in untreated controls.

#### In vivo studies

Tumor organoids were collected, incubated with TripLE at 37°C for 10 min, dissociated into single cells, resuspended in tumor organoid medium, filtered with 70µm nilon filters (Corning) and mixed in a in complete mouse media/BME mixture (1:1). KB1P4.N1 and KB1P4.R1 organoid suspensions contained a total of 20.000 and 10.000 cells, respectively, per 40 µl of media/BME mixture, and were injected in the fourth right mammary fat pad of wild-type FVB/N mice. Mammary tumor size was determined by caliper measurements (length and width in millimeters), and tumor volume (in mm^3^) was calculated by using the following formula: 0.5 × length × width^2^. Upon tumor outgrowth to approximately 75 mm^3^, in mice injected with N1 organoids, and 40 mm^3^, in mice injected with R1 organoids, mice were treated with vehicle, or olaparib (50 mg/kg, mice injected with N1 organoids; 100 mg/kg, mice injected with R1 organoids) intraperitoneally for 28 consecutive days. Animals were sacrificed with CO_2_ when the tumor volume reached 1,500 mm^3^.

#### Immunohistochemistry Analysis

Five-µm tissue sections were cut from formalin-fixed, paraffin-embedded tissue blocks from a cohort of 86 TNBC (Gogola *et al*., 2018) and 51 human serous ovarian carcinomas (Moudry et al. 2016) and mounted on Super Frost Plus slides (Menzel-Glaser, Braunschweig, Germany), baked at 60°C for 60 min, deparaffinized, and rehydrated through graded alcohol rinses. Heat induced antigen retrieval was performed by immersing the slides in citrate pH 6.0 buffer and heating them in a 750 W microwave oven for 15 min. The sections were then stained with primary antibody anti-LIG3 (1: 250, Sigma-Aldrich, #HPA006723) overnight in a cold-room, followed by subsequent processing by the indirect streptavidin-biotin-peroxidase method using the Vectastain Elite kit (Vector Laboratories, Burlingame, CA, USA) and nickel-sulphate-based chromogen enhancement detection as previously described (Bartkova *et al*., 2005), without nuclear counterstaining. For negative controls, sections were incubated with non-immune sera. The results were evaluated by two experienced researchers, including a senior oncopathologist, and the data expressed as percentage of positive tumor cells within each lesion, while recording frequencies of cases with LIG3 overabundant (LIG3-high) or lost (LIG3-low) staining in 10-20% and in excess of 20% of the tumor cells (see Figure 6G for examples of staining patterns). Cases with over 90% of cancer cells showing a staining intensity comparable with surrounding stromal cells on the same section (internal control) were regarded as displaying a normal pattern of LIG 3 expression.

### QUANTIFICATION AND STATISTICAL ANALYSIS

Statistical analysis was performed using Prism (GraphPad Software), unless stated in the figure legend. In all cases: ns, non-significant, * p <0.05, ** p <0.01, *** p <0.001, **** p <0.0001. For long-term clonogenic and short-term cytotoxicity assays, qRT-PCR and analysis of metaphase spreads, two-tailed unpaired t test was used. For immunofluorescence, unpaired t test was used. For DNA fiber analysis and PLA, group comparisons were performed with Mann–Whitney U test. Analysis of EM was carried out using two-way ANOVA. For survival analysis, data are presented as Kaplan-Meier curves and the p values were computed using Log-Rank (Mantel Cox) statistics.

### DATA AND SOFTWARE AVAILABILITY

This study did not generate/analyze datasets/code.

## SUPPLEMENTAL ITEM TITLES

**Figure S1.** Depletion of LIG3 Increases Sensitivity to PARPi in HR-Negative and HR-Restored Cells. Related to Figure 1.

**Figure S2.** Lethality Observed in LIG3-Depleted Cells is Dependent on BRCA1 Loss. Related to Figure 1.

**Figure S3.** PARP1 Trapping Contributes to PARPi Toxicity in LIG3-Depleted cells.

**Figure S4.** Resistance to PARPi in 53BP1-Deficient KB1P Cells is Mediated by Nuclear LIG3. Related to Figure 2.

**Figure S5.** LIG3 is Required at Replication Forks in BRCA1-Deficient Cells Treated with PARPi. Related to Figure 3.

**Figure S6.** LIG3 Depletion Reverts PARPi Resistance by Increasing Post-replicative MRE11- Mediated ssDNA Gaps. Related to Figures 4 and 5.

**Figure S7.** LIG3 Depletion Does Not Result in DSB Formation. Related to Figure 5.

**Table S1.** Gene p value for T0, untreated and treated conditions for both screens, analyzed by MAGeCK and DESeq2. Related to Figure 1.

**Table S2.** Oligonucleotides used in this study. Related to STAR METHODS key resource table.

