## Supplemental Figures for "Loss of Nuclear DNA Ligase III Reverts PARP Inhibitor Resistance in BRCA1/53BP1 Double-deficient Cells by Exposing ssDNA Gaps"

FIGURE S1

A

| Cell line | Genotype | Cell line | Genotype |
|---|---|---|---|
| ORG-KB1P.R | *Brca1*$^{-/-}$;*Trp53*$^{-/-}$;*Trp53bp1*$^{-/-}$ | ES-P.R | *Brca1*$^{SCo/-}$;*Trp53*$^{-/-}$;*Trp53bp1*$^{-/-}$ |
| KB1P.R | | ES-B1P.R | *Brca1*$^{-/-}$;*Trp53*$^{-/-}$;*Trp53bp1*$^{-/-}$ |
| ORG-KB1P.S | *Brca1*$^{-/-}$;*Trp53*$^{-/-}$ | ES-P | *Brca1*$^{SCo/-}$;*Trp53*$^{-/-}$ |
| KB1P.S | | ES-B1P.S | *Brca1*$^{-/-}$;*Trp53*$^{-/-}$ |
| ORG-KP | *Trp53*$^{-/-}$ | RPE1-B1P.R | *BRCA1*$^{-/-}$;*TP53*$^{-/-}$;*TP53BP1*$^{-/-}$ |
| KP | | RPE1-B1P.S | *BRCA1*$^{-/-}$;*TP53*$^{-/-}$ |
| KB2P | *Brca2*$^{-/-}$;*Trp53*$^{-/-}$ | RPE1-P | *TP53*$^{-/-}$ |

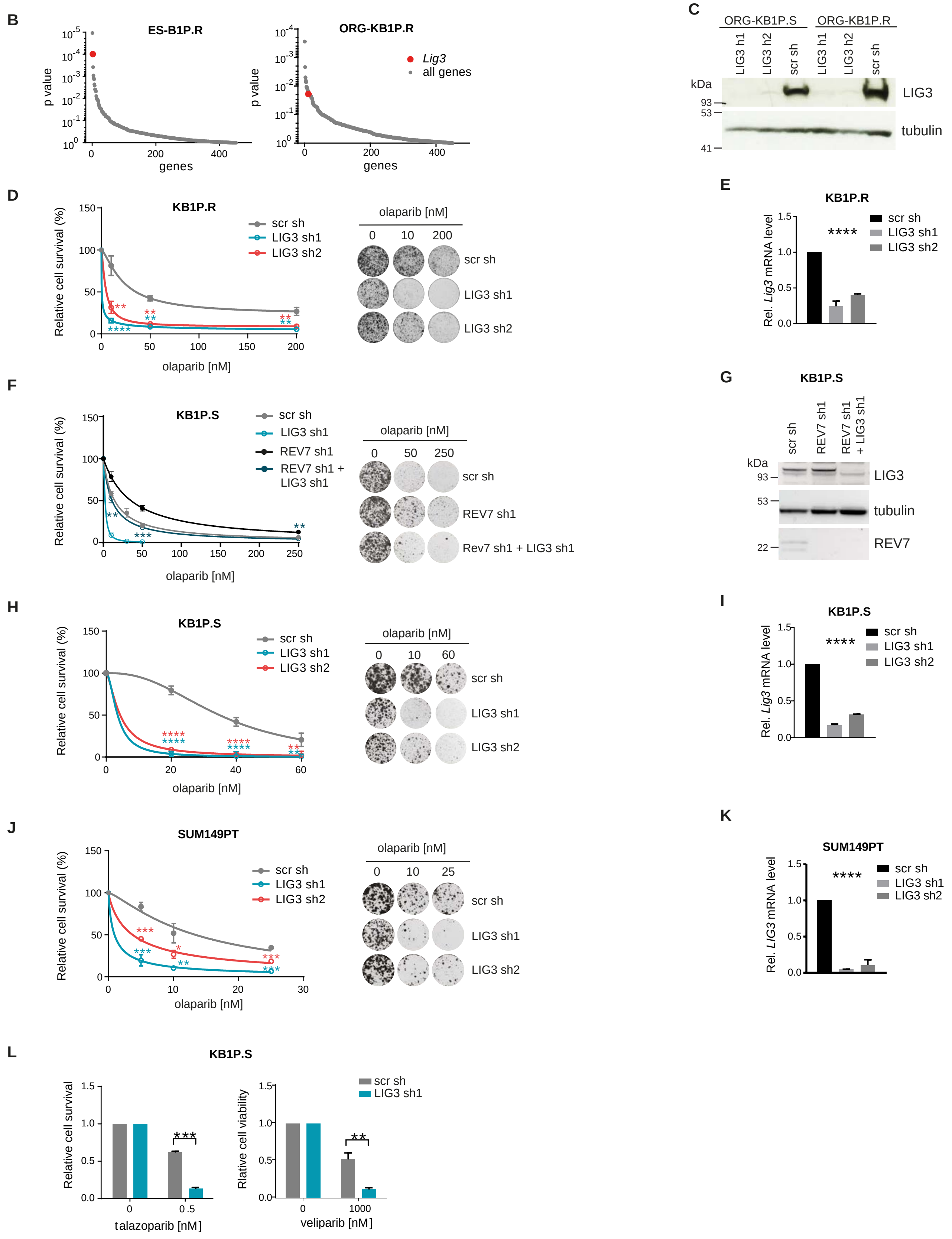

**Depletion of LIG3 Increases Sensitivity to PARPi in HR-Negative and HR-Restored Cells. Related to Figure 1.**

**(A)** Summary table of cell line abbreviations and respective genotypes.
**(B)** Distribution of the one-sided p value (gene dropout) for all genes targeted by the shRNA-based library in mESCs (left) and in organoids (right), by MAGeCK.
**(C)** Western blot analysis of LIG3 in ORG-KB1P.S and ORG-KB1P.R organoids, transduced with shRNA targeting LIG3.
**(D)** Quantification (left) and representative images (right) of long-term clonogenic assay with KB1P.R cells treated with olaparib or left untreated.
**(E)** RT-qPCR analysis of *Lig3* expression levels in KB1P.R cells expressing the indicated shRNAs.
**(F)** Quantification (left) and representative images (right) of long-term clonogenic assay with KB1P.S cells, treated with olaparib or left untreated.
**(G)** Western blot analysis of LIG3 and REV7 in KB1P.S cells expressing indicated shRNAs.
**(H)** Quantification (left) and representative images (right) of long-term clonogenic assay with KB1P.S cells, treated with olaparib or left untreated.
**(I)** RT-qPCR analysis of *Lig3* expression levels in KB1P.S cells expressing the indicated shRNAs.
**(J)** Quantification (left) and representative images (right) of long-term clonogenic assay with SUM149PT cells, treated with olaparib or left untreated.
**(K)** RT-qPCR analysis of *LIG3* expression levels in SUM149PT cells expressing the indicated shRNAs.
**(L)** Quantification of long-term clonogenic assay with KB1P.S cells, treated with the PARPi talazoparib (left) and veliparib (right).
Data are represented as mean ± SD. * $p<0.05$, ** $p<0.01$, *** $p<0.001$, **** $p<0.0001$, n.s., not significant; two-tailed t test.

FIGURE S2

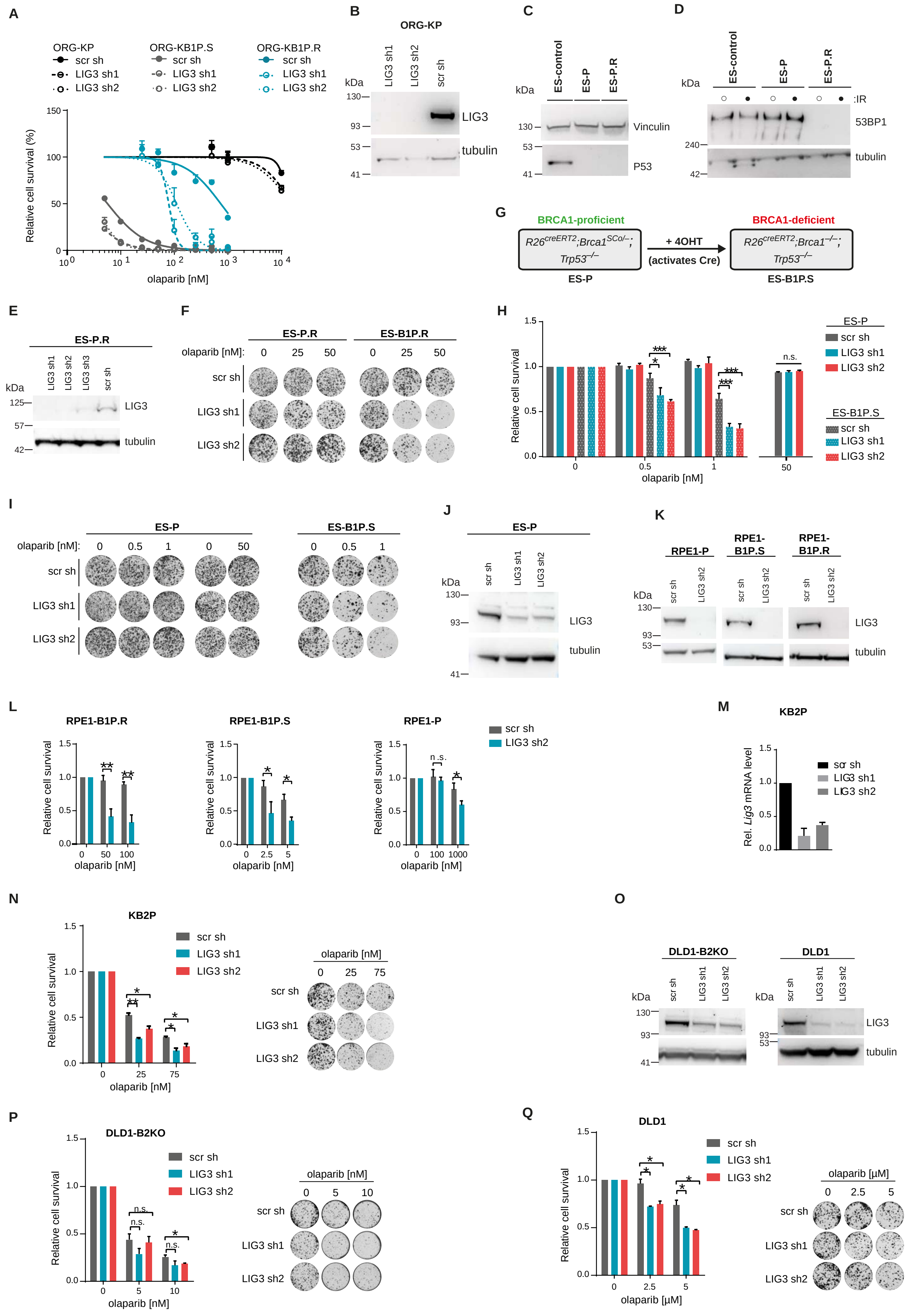
A
ORG-KP
scr sh
LIG3 sh1
LIG3 sh2
ORG-KB1P.S
ORG-KB1P.R
Relative cell survival (%)
olaparib [nM]
B
ORG-KP
LIG3
tubulin
C
ES-control
ES-P
ES-P.R
Vinculin
P53
D
53BP1
:IR
G
BRCA1-proficient
BRCA1-deficient
+ 4OHT
(activates Cre)
ES-P
ES-B1P.S
E
ES-P.R
F
ES-B1P.R
olaparib [nM]:
H
Relative cell survival
ES-B1P.S
I
J
K
RPE1-P
RPE1-B1P.S
RPE1-B1P.R
L
M
KB2P
Rel. Lig3 mRNA level
N
O
DLD1-B2KO
DLD1
P
Q
olaparib [μM]

**Lethality Observed in LIG3-Depleted Cells is Dependent on BRCA1 Loss. Related to Figure 1.**

**(A)** Quantification of long-term clonogenic assay with ORG-KB1P.R, ORG-KB1P.S, and ORG-KP organoids treated with olaparib.
**(B)** Western blot analysis of LIG3 in ORG-KP organoids, transduced with shRNA targeting LIG3.
**(C and D)** Western blot analysis of P53 **(C)** and 53BP1 **(D)** in ES-P.R and in ES-P mESCs.
**(E)** Western blot analysis of LIG3 in ES-P.R mESCs, transduced with shRNA targeting LIG3.
**(F)** Representative images of long-term clonogenic assay with ES.P.R and ES-B1P.R mESCs treated with olaparib or left untreated.
**(G)** Schematic representation of the *Brca1* selectable conditional allele in $R26^{creERT2}$;$Brca1^{SCo/–}$;$Trp53^{–/–}$ (ES-P). Incubation of these cells with 4-hydroxytamoxifen (4OHT) induces a CreERT2 recombinase fusion protein, resulting in $R26^{creERT2}$;$Brca1^{–/–}$;$Trp53^{–/–}$ (ES-B1P.R) cells lacking BRCA1 protein expression.
**(H and I)** Quantification **(H)** and representative images **(I)** of long-term clonogenic assay with ES-P and ES-B1P.S mESCs treated with olaparib or left untreated.
**(J)** Western blot analysis of LIG3 in ES-P mESCs, transduced with shRNA targeting LIG3.
**(K)** Western blot analysis of LIG3 in RPE1-P, RPE1-B1P.S and RPE1-B1P.R cells transduced with shRNA targeting LIG3.
**(L)** Quantification of long-term clonogenic assay with RPE1-P, RPE1-B1P.S and RPE1-B1P.R cells treated with olaparib or left untreated.
**(M)** RT-qPCR analysis of *Lig3* expression levels in KB1P.R cells expressing indicated shRNAs.
**(N)** Quantification (left) and representative images (right) of long-term clonogenic assay with KB2P cells treated with olaparib or left untreated.
**(O)** Western blot analysis of BRCA2-deficient DLD1-B2KO cells and parental DLD1 cells transduced with LIG3-targeting shRNAs.
**(Pand Q)** Quantification (left) and representative images (right) of long-term clonogenic assay with DLD1-B2KO **(P)** and parental DLD1 cells **(Q)** treated with olaparib or left untreated.
Data are represented as mean ± SD. * $p<0.05$, ** $p<0.01$, *** $p<0.001$, **** $p<0.0001$, n.s., not significant; two-tailed t test.

FIGURE S3

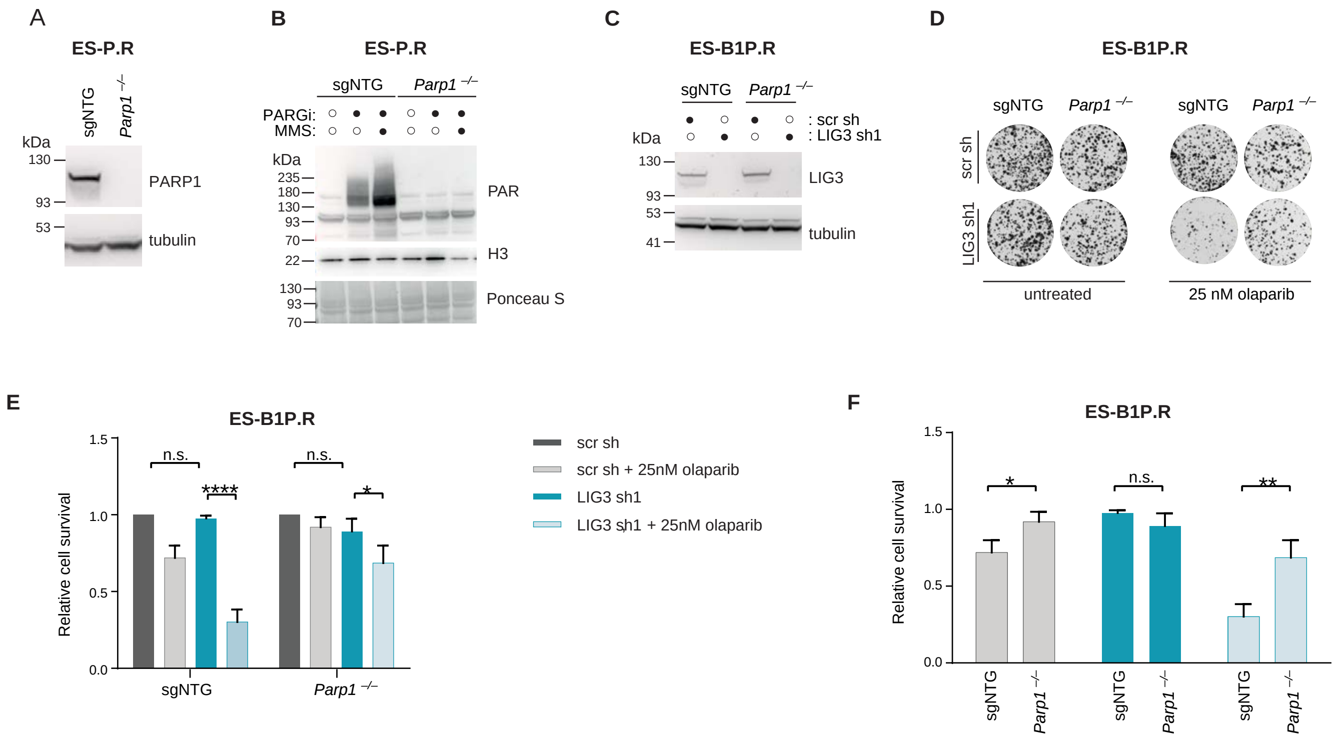


**PARP1 Trapping Contributes to PARPi Toxicity in LIG3-Depleted cells.**

**(A)** Western blot analysis of PARP1 in ES-P.R cells transduced with non-targeting single-guide RNA (ES-P.R sgNTG) and in ES-P.R *Parp1*$^{-/-}$ cells.
**(B)** Western blot analysis of PAR in ES-P.R sgNTG and in ES-P.R *Parp1*$^{-/-}$ cells, left untreated, or treated with PARGi (PDDX-001) and/or 0.01% MMS for 30 min.
**(C)** Western blot analysis of LIG3 in ES-P.R sgNTG and ES-P.R *Parp1*$^{-/-}$ cells, transduced with shRNA targeting LIG3.
**(D-F)** Representative images **(D)** and quantification **(E,F)** of long-term clonogenic assay in ES-B1P.R *Parp1*$^{-/-}$ cells treated with olaparib and upon shRNA-mediated depletion of LIG3. Values were normalized to untreated scr sh for each line.
Data are represented as mean ± SD. * $p<0.05$, **$p<0.01$, ***$p<0.001$, ****$p<0.0001$, n.s., not significant; two-tailed t test.

FIGURE S4

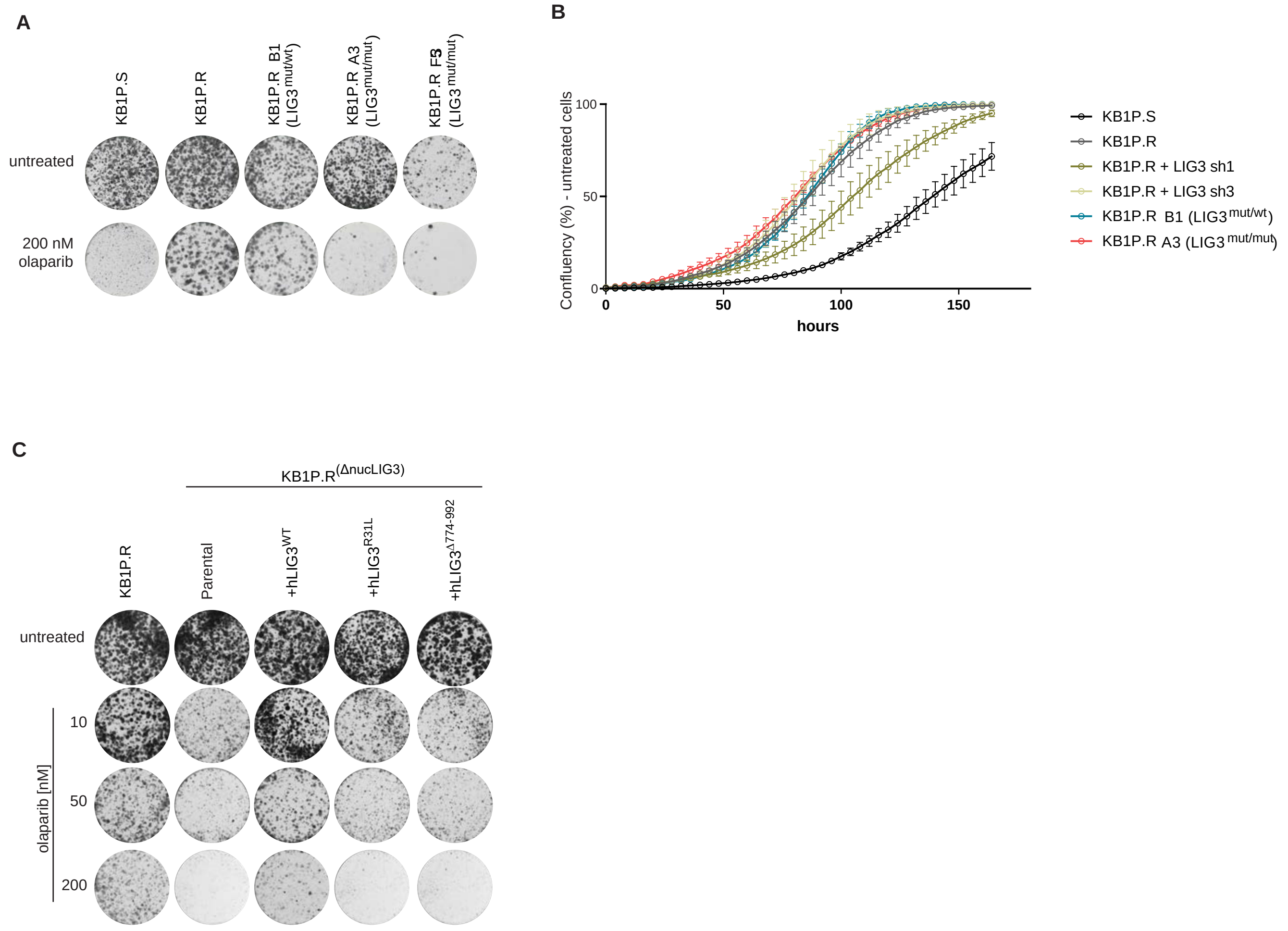


**Resistance to PARPi in 53BP1-Deficient KB1P Cells is Mediated by Nuclear LIG3. Related to Figure 2.**

**(A)** Representative images of long-term clonogenic assay with KB1P.S, KB1P.R, KB1P.R(LIG3$^{mut/wt}$) B1, KB1P.R(LIG3$^{mut/mut}$) A3 and KB1P.R(LIG3$^{mut/mut}$) F5 cells, treated with olaparib or left untreated.

**(B)** Quantification of proliferation assays in KB1P.S, KB1P.R, KB1P.R transduced with shRNAs targeting LIG3, KB1P.R(LIG3$^{mut/mut}$) A3 and KB1P.R(LIG3$^{mut/wt}$) B1. Cell confluency was measured every 4h with IncuCyte.

**(C)** Representative images of long-term clonogenic assay with KB1P.R and KB1P.R$^{(\Delta nucLIG3)}$ nuclear LIG3 mutant cells (for which we selected KB1P.R(LIG3$^{mut/mut}$) clone A3), treated with olaparib or untreated. Expression of indicated LIG3 constructs was induced with Doxycycline 2 days before the assay and maintained for the duration of the assay.

FIGURE S5

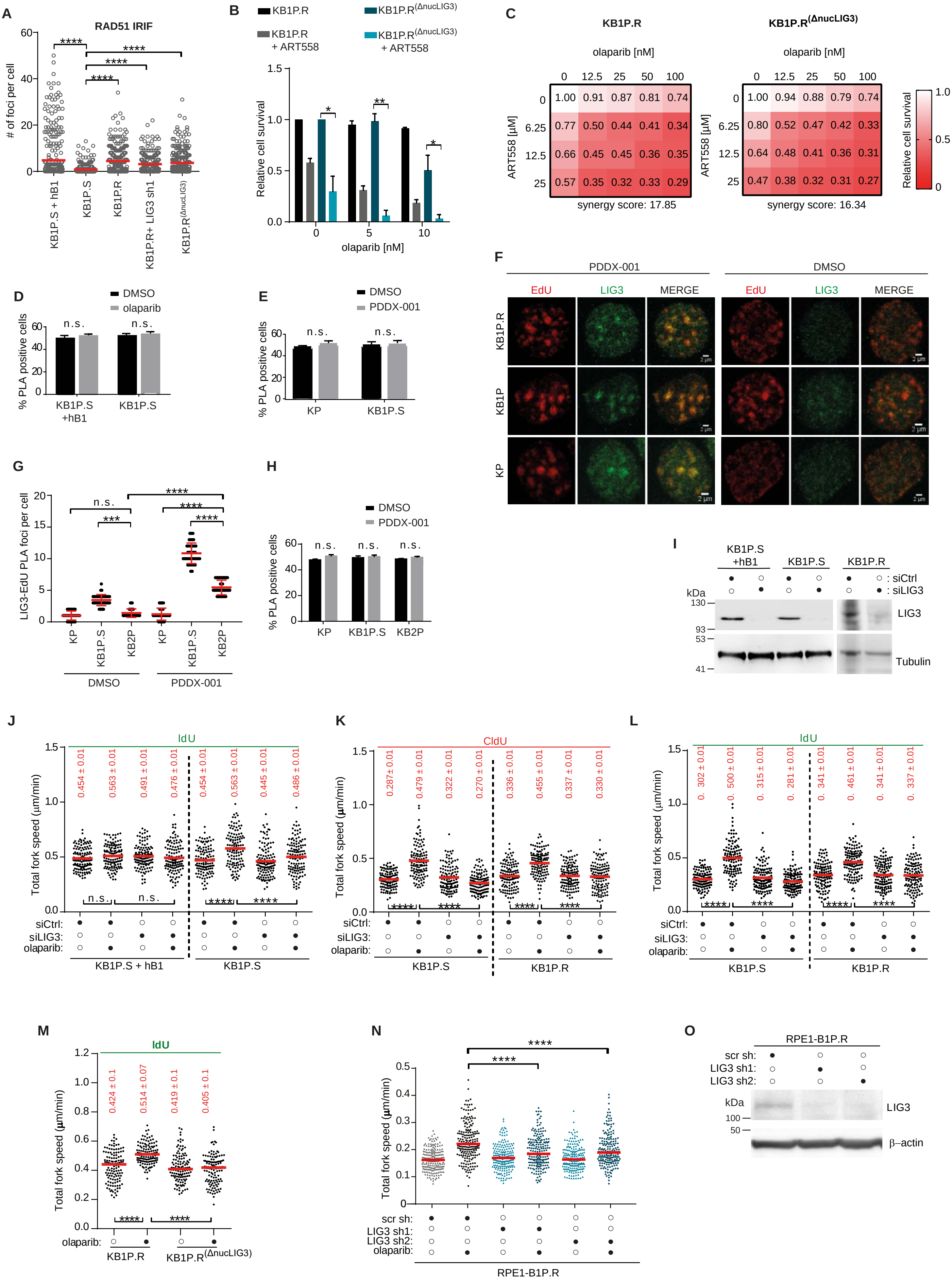
A
RAD51 IRIF
### of foci per cell
KB1P.S + hB1
KB1P.S
KB1P.R
KB1P.R+ LIG3 sh1
KB1P.R(ΔnucLIG3)
B
KB1P.R
KB1P.R + ART558
KB1P.R(ΔnucLIG3)
KB1P.R(ΔnucLIG3) + ART558
Relative cell survival
olaparib [nM]
C
KB1P.R
KB1P.R(ΔnucLIG3)
olaparib [nM]
ART558 [μM]
synergy score: 17.85
synergy score: 16.34
Relative cell survival
D
DMSO
olaparib
% PLA positive cells
KB1P.S +hB1
KB1P.S
E
DMSO
PDDX-001
% PLA positive cells
KP
KB1P.S
F
PDDX-001
DMSO
EdU
LIG3
MERGE
KB1P.R
KB1P
KP
G
LIG3-EdU PLA foci per cell
KP
KB1P.S
KB2P
DMSO
PDDX-001
H
DMSO
PDDX-001
% PLA positive cells
KP
KB1P.S
KB2P
I
KB1P.S +hB1
KB1P.S
KB1P.R
: siCtrl
: siLIG3
kDa
LIG3
Tubulin
J
IdU
Total fork speed (μm/min)
0.454 ± 0.01
0.563 ± 0.01
0.491 ± 0.01
0.476 ± 0.01
0.454 ± 0.01
0.563 ± 0.01
0.445 ± 0.01
0.486 ± 0.01
siCtrl:
siLIG3:
olaparib:
KB1P.S + hB1
KB1P.S
K
CldU
Total fork speed (μm/min)
0.287± 0.01
0.479 ± 0.01
0.322 ± 0.01
0.270 ± 0.01
0.336 ± 0.01
0.455 ± 0.01
0.337 ± 0.01
0.330 ± 0.01
siCtrl:
siLIG3:
olaparib:
KB1P.S
KB1P.R
L
IdU
Total fork speed (μm/min)
siCtrl:
siLIG3:
olaparib:
KB1P.S
KB1P.R
M
IdU
Total fork speed (μm/min)
0.424 ± 0.1
0.514 ± 0.07
0.419 ± 0.1
0.405 ± 0.1
olaparib:
KB1P.R
KB1P.R(ΔnucLIG3)
N
Total fork speed (μm/min)
scr sh:
LIG3 sh1:
LIG3 sh2:
olaparib:
RPE1-B1P.R
O
RPE1-B1P.R
scr sh:
LIG3 sh1:
LIG3 sh2:
kDa
LIG3
β–actin

**LIG3 is Required at Replication Forks in BRCA1-Deficient Cells Treated with PARPi. Related to Figure 3.**

**(A)** Quantification of RAD51 IRIF after irradiation with 10Gy and 3 hr recovery, in KP, KB1P.S, KB1P.R, KB1P.R after shRNA-mediated LIG3 depletion, and nuclear LIG3 mutant KB1P.R$^{(\Delta nucLIG3)}$ cells. Data are represented as mean. ****p<0.0001, Unpaired T test.

**(B)** Quantification of long-term clonogenic assay in KB1P.R and nuclear LIG3 mutant KB1P.R$^{(\Delta nucLIG3)}$ cells, left untreated or treated with olaparib and/or 25μM POLθ inhibitor ART558. Treatment with olaparib was carried out at concentrations not toxic to KB1P.R$^{(\Delta nucLIG3)}$ cells so epistasis or absence of it could be observed. Data are represented as mean ± SD. * p<0.05, **p<0.01, n.s., not significant; two-tailed t test.

**(C)** Quantification of short-term cytotoxicity assay upon combination treatment with olaparib and POLθ inhibitor ART558, at the indicated concentrations, in KB1P.R and KB1P.R$^{(\Delta nucLIG3)}$. Treatment with olaparib was carried out at concentrations not toxic to KB1P.R$^{(\Delta nucLIG3)}$ cells so epistasis or absence of it could be observed. Synergy scores were calculated based on Bliss reference model using SynergyFinder.

**(D)** Percentage of LIG3-EdU proximity ligation assay (PLA) positive cells corresponding to Figure 4A.

**(E)** Percentage of LIG3-EdU PLA positive cells corresponding to Figure 4B.

**(F)** Immunostaining of LIG3 in detergent-pre-extracted KB1P.R, KB1P.S and KP cells, incubated for 1 hr with 20μM EdU in the absence or presence of PARG inhibior PDDX-001.

**(G)** Quantification of LIG3-EdU PLA foci in KB2P cells incubated for 10 min with 20μM EdU, in the absence or presence of PDDX-001. ± SD,***p<0.001 ****p<0.0001; n.s., not significant; Mann–Whitney U test.

**(H)** Percentage of LIG3-EdU PLA positive cells in (F).

**(I)** Western blot analysis of LIG3 in KB1P.S+hB1, KB1P.S and KB1P.R cells transfected with siRNA targeting LIG3, used for DNA fiber assays.

**(J)** Quantification of IdU tracks in KB1P.S+hB1 and KB1P.S cells, following the indicated treatments. Data are represented as mean. ****p<0.0001, n.s., not significant; Mann–Whitney U test.

**(K and L)** Quantification of CldU **(J)** and IdU tracks **(K)** in KB1P.S and KB1P.R cells, following the indicated treatments. Data are represented as mean. n.s., not significant, ****p<0.0001, Mann–Whitney U test.

**(M)** Quantification of IdU tracks in nuclear LIG3- mutant KB1P.R$^{(\Delta nucLIG3)}$ cells, following the indicated treatments. Data are represented as mean. ****p<0.0001, n.s., not significant; Mann–Whitney U test.

**(N)** Quantification of fork speed in RPE1-B1P.R cells transfected with shRNAs targeting LIG3, following the indicated treatments. Data are represented as mean. ****p<0.0001, Mann–Whitney U test.

**(O)** Western blot analysis of RPE1-B1P.R cells after shRNA-mediated depletion of LIG3.

FIGURE S6

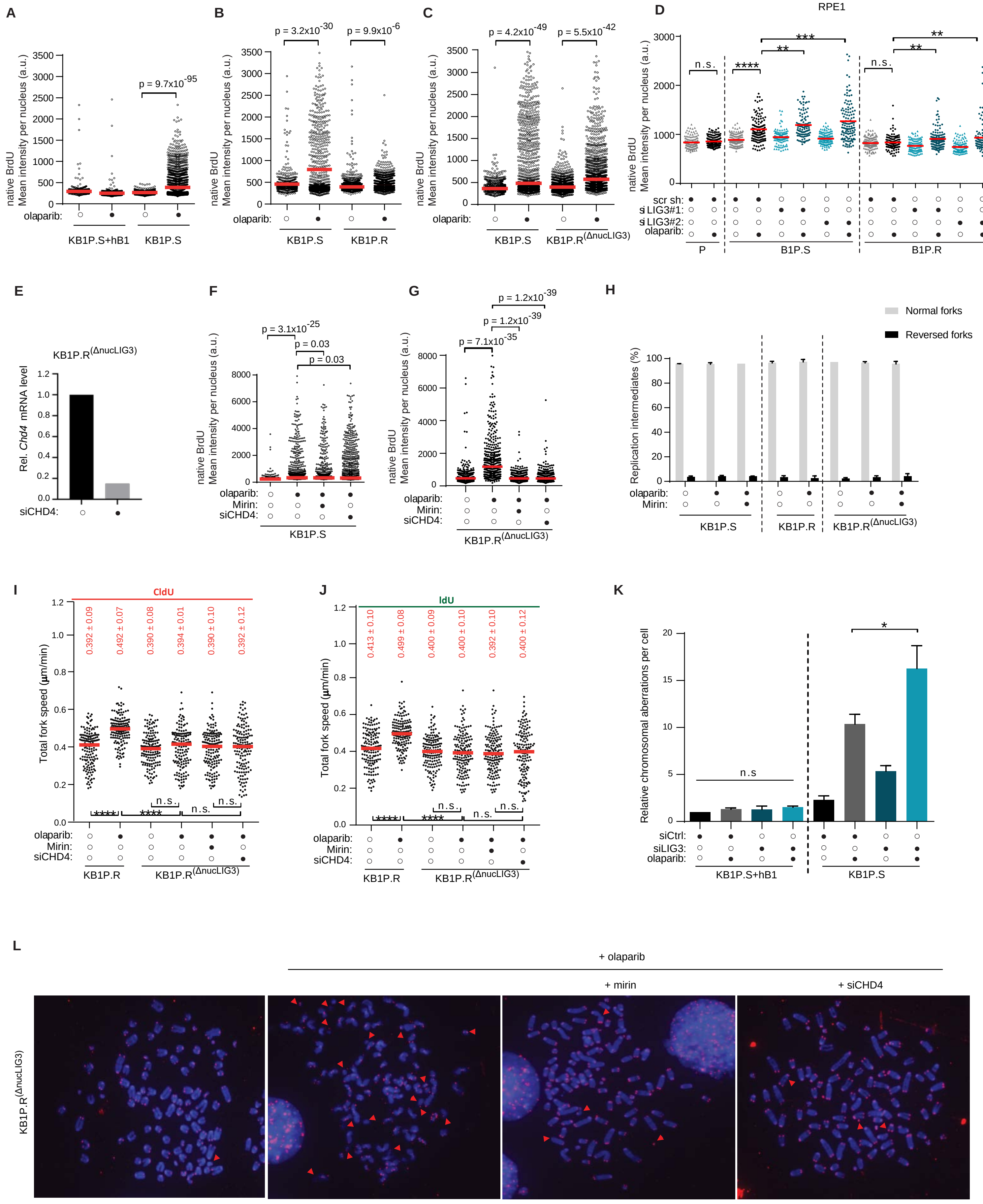

A
native BrdU Mean intensity per nucleus (a.u.)
p = 9.7x10^-95
olaparib:
KB1P.S+hB1
KB1P.S
B
p = 3.2x10^-30
p = 9.9x10^-6
native BrdU Mean intensity per nucleus (a.u.)
olaparib:
KB1P.S
KB1P.R
C
p = 4.2x10^-49
p = 5.5x10^-42
native BrdU Mean intensity per nucleus (a.u.)
olaparib:
KB1P.S
KB1P.R(ΔnucLIG3)
D
RPE1
native BrdU Mean intensity per nucleus (a.u.)
n.s.
****
**
***
n.s.
**
**
scr sh:
si LIG3#1:
si LIG3#2:
olaparib:
P
B1P.S
B1P.R
E
KB1P.R(ΔnucLIG3)
Rel. Chd4 mRNA level
siCHD4:
F
p = 3.1x10^-25
p = 0.03
p = 0.03
native BrdU Mean intensity per nucleus (a.u.)
olaparib:
Mirin:
siCHD4:
KB1P.S
G
p = 1.2x10^-39
p = 1.2x10^-39
p = 7.1x10^-35
native BrdU Mean intensity per nucleus (a.u.)
olaparib:
Mirin:
siCHD4:
KB1P.R(ΔnucLIG3)
H
Normal forks
Reversed forks
Replication intermediates (%)
olaparib:
Mirin:
KB1P.S
KB1P.R
KB1P.R(ΔnucLIG3)
I
CldU
Total fork speed (µm/min)
0.392 ± 0.09
0.492 ± 0.07
0.390 ± 0.08
0.394 ± 0.01
0.390 ± 0.10
0.392 ± 0.12
****
****
n.s.
n.s.
n.s.
olaparib:
Mirin:
siCHD4:
KB1P.R
KB1P.R(ΔnucLIG3)
J
IdU
Total fork speed (µm/min)
0.413 ± 0.10
0.499 ± 0.08
0.400 ± 0.09
0.400 ± 0.10
0.392 ± 0.10
0.400 ± 0.12
****
****
n.s.
n.s.
n.s.
olaparib:
Mirin:
siCHD4:
KB1P.R
KB1P.R(ΔnucLIG3)
K
Relative chromosomal aberrations per cell
n.s
*
siCtrl:
siLIG3:
olaparib:
KB1P.S+hB1
KB1P.S
L
+ olaparib
+ mirin
+ siCHD4
KB1P.R(ΔnucLIG3)

**LIG3 Depletion Reverts PARPi Resistance by Increasing Post-replicative MRE11-Mediated ssDNA Gaps. Related to Figures 4 and 5.**

**(A-C)** Dot plot of native BrdU mean intensity per nucleus in shown in Figure 4B **(A),** in Figure 4C **(B)** and in Figure 4D **(C)**. Data are represented as mean. Unpaired t test, p value was calculated using R.
**(D)** Quantification of immunofluorescense analysis of ssDNA gaps in RPE1 cells as shown in 4A. * p<0.05, **p<0.01, ***p<0.001, ****p<0.0001, n.s., not significant; unpaired t test.
**(E)** RT-qPCR analysis of *Chd4* expression levels in nuclear LIG3 mutant KB1P.R$^{(\Delta nucLIG3)}$ cell line transfected siRNA targeting CHD4.
**(F and G)** Dot plot of native BrdU mean intensity per nucleus shown in Figure 5D. Unpaired t test, p value was calculated using R.
**(H)** Quantification of normal and reversed forks in KB1P.S, KB1P.R and KB1P.R$^{(\Delta nucLIG3)}$ cells. Data were acquired by electron microscopy.
**(I and J)** Quantification of CldU **(I)** and IdU tracks **(J)** in KB1P.R and KB1P.R$^{(\Delta nucLIG3)}$ cells, following the indicated treatments. KB1P.R$^{(\Delta nucLIG3)}$ cells were additionally treated with 25μM mirin or transfected with siRNA targeting CHD4, 48 hr prior to treatment with olaparib. Data are represented as mean. ****p<0.0001, n.s., not significant; Mann–Whitney U test.
**(K)** Quantification of chromosomal aberrations in LIG3-proficient and LIG3-depleted KB1P.S+hB1 and KB1P.S cells following 2 hr treatment with 0.5μM olaparib and recovery for 6 hr.
**(L)** Representative images of chromosomal aberrations in Figure 5F and 5G. Telomeres are labeled in red. Red arrowheads indicate chromosomal aberrations.

FIGURE S7

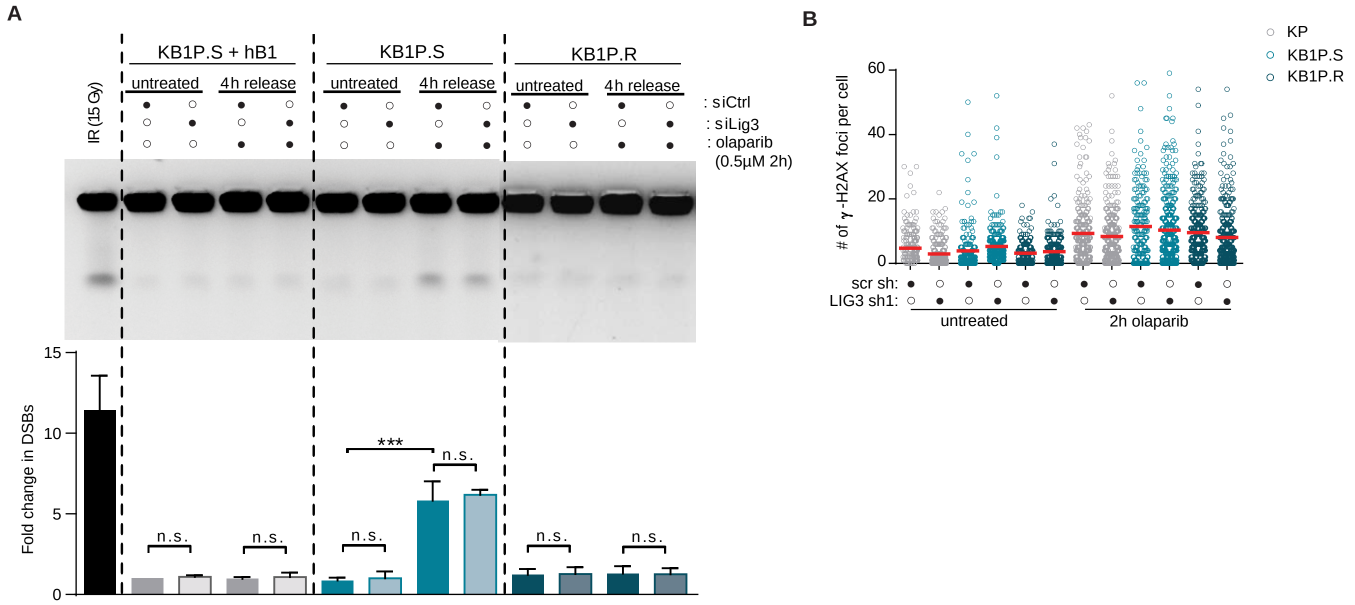


**LIG3 Depletion Does Not Result in DSB Formation. Related to Figure 6.**

**(A)** Representative image (top) and quantification (bottom) of pulsed-field gel electrophoresis (PFGE) analysis of DSBs in LIG3-proficient and LIG3-depleted KB1P.S+hB1, KB1P.S and KB1P.R cells, treated with 0.5µM olaparib for 2 hr and released for 4hr or left untreated. Data are represented as mean ± SD. ***$p<0.001$, n.s., not significant, two-tailed t test.

**(B)** γ-H2AX foci formation in LIG3-proficient and LIG3-depleted KP, KB1P.S and KB1P.R cells treated with 0.5µM olaparib for 2 hr or left untreated.

.
