## Supplementary material for "Loss of Nuclear DNA Ligase III Reverts PARP Inhibitor Resistance in BRCA1/53BP1 Double-deficient Cells by Exposing ssDNA Gaps": Table S2

| OLIGONUCLEOTIDE | EXPERIMENT | SEQUENCE |
|---|---|---|
| pLKO.1-scrambled shRNA (lentiviral) | shRNA-mediated knockdown | CCTAAGGTTAAGTCGCCCTCG |
| pLKO.1- *Lig3* shRNA #1 (mouse, lentiviral) | shRNA-mediated knockdown | CCAGACTTCAAACGTCTCAAA |
| pLKO.1- *Lig3* shRNA #2 (mouse, lentiviral) | shRNA-mediated knockdown | CGTGTCAGAGACGATCAGAAT |
| pLKO.1- *Rev7* shRNA (mouse, lentiviral) | shRNA-mediated knockdown | CCCGGAGCTGAATCAGTATAT |
| pLKO.1- *LIG3* shRNA #1 (human, lentiviral) | shRNA-mediated knockdown | GCCCACTTTAAGGACTACATT |
| pLKO.1- *LIG3* shRNA #2 (human, lentiviral) | shRNA-mediated knockdown | CCGGATCATGTTCTCAGAAAT |
| NT (non-targeting) gRNA | CRISPR/Cas9 genome editing | TGATTGGGGGTCGTTCGCCA |
| mouse *Trp53* sgRNA | CRISPR/Cas9 genome editing | GAAGTCACAGCACATGACGG |
| mouse *Trp53bp1* sgRNA | CRISPR/Cas9 genome editing | TACCGGGCTGTACTGTAACA |
| mouse *Parp1* sgRNA | CRISPR/Cas9 genome editing | CGAGTGGAGTACGCGAAGAG |
| mouse *Lig3* sgRNA - point mutation | CRISPR/Cas9 genome editing | CTGTACTGGCCCCTGTGCGA |
| ssODN - point mutation template forward | Homology-mediated CRISPR/Cas9 genome editing | GCCACCCACCTTACTTTCTGGCCAGGGTCGCATG TGGGACTCTGTACTGGCCCCTGTGCGCTCGCAG AGCAGCGGTTCTGTGTGGACTATGCCAAGCGGG GCACAGCTGGATGCAAGAAA |
| ssODN - point mutation template reverse | Homology-mediated CRISPR/Cas9 genome editing | TTTCTTGCATCCAGCTGTGCCCCGCTTGGCATAG TCCACACAGAACCGCTGCTCTGCGAGCGCACAG GGGCCAGTACAGAGTCCCACATGCGACCCTGGC CAGAAAGTAAGGTGGGTGGC |
| point mutation template control for TIDE forward | TIDE analysis | ACTGGCCCCTGTGCGCTCGCA GAGCAGCGGTTCTGTGTGGAC |
| point mutation template control for TIDE reverse | TIDE analysis | GTCCACACAGAACCGCTGCTC TGCGAGCGCACAGGGGCCAGT |
| mouse *Trp53* sgRNA forward primer | TIDE analysis | CCCACCTTGACACCTGATCG |
| mouse *Trp53* sgRNA reverse primer | TIDE analysis | CCACCCGGATAAGATGCTGG |
| mouse *Trp53bp1* sgRNA forward primer | TIDE analysis | GAGAGCGCACGCACAGTAAG |
| mouse *Trp53bp1* sgRNA reverse primer | TIDE analysis | TGGGCTGGCTCTGATACTTTG |
| mouse *Parp1* sgRNA forward primer | TIDE analysis | AACCGACAAAAGGGGTGGCG |
| mouse *Parp1* sgRNA reverse primer | TIDE analysis | GCAGGGTAAGCGCAATGTCC |
| mouse *Lig3* forward primer | RT-qPCR | GAAATTGCTGCGGCACCATTA |
| mouse *Lig3* reverse primer | RT-qPCR | AGCCATCATTGTAGTTGACCTG |
| human *HPRT* forward primer | RT-qPCR | GAAGAGCTATTGTAATGACC |
| human *HPRT* reverse primer | RT-qPCR | GCGACCTTGACCATCTTTG |
| mouse *Rev7* forward primer | RT-qPCR | ACACTCCACTGCGTCAAACC |
| mouse *Rev7* reverse primer | RT-qPCR | AAAGACAAACTTCTCCACTGGGC |
| mouse *Lig3* forward primer | RT-qPCR | TTACCAGTACCAATCCTCGGAA |
| mouse *Lig3* reverse primer | RT-qPCR | ACAATCTTTGTGCTTAGGGTCAC |
| mouse *Rps20* forward primer | RT-qPCR | TGTGCGGACTTGATCAGAGG |
| mouse *Rps20* reverse primer | RT-qPCR | GGTCTTGGAACCTTCACCACA |
| human *LIG3* forward primer | RT-qPCR | GCCGGAGAGGCAGCTATATG |
| human *LIG3* reverse primer | RT-qPCR | GGCAACAGTTCTTTTCGGCTG |
